# VDisk: A Microfluidic Cartridge for Automated and Reproducible Label-Free Isolation of Extracellular Vesicles from Plasma

**DOI:** 10.1101/2025.09.04.674190

**Authors:** Ehsan Mahmodi Arjmand, Gustav Grether, Gonzalo Bustos-Quevedo, John Atanga, Pablo Sánchez-Martín, Jan Van Deun, Tobias Hutzenlaub, Irina Nazarenko, Nils Paust, Jan Lüddecke

**Affiliations:** Hahn-Schickard, Freiburg, Germany; Laboratory for MEMS Applications, IMTEK-Department of Microsystems Engineering, University of Freiburg, Freiburg, Germany; Institute for Infection Prevention and Control, Medical Center and Faculty of Medicine, University of Freiburg, Freiburg, Germany; Faculty of Biology, University of Freiburg, Germany; German Cancer Consortium (DKTK), Partner Site Freiburg, and German Cancer Research Center (DKFZ), Heidelberg, Germany; PICO BioScience GmbH, Freiburg, Germany; Department of Dermatology, University Hospital Erlangen, Erlangen, Germany

**Keywords:** Extracellular vesicles, liquid biopsy, plasma, automation, chromatography, reproducibility, centrifugal microfluidics

## Abstract

Blood-derived extracellular vesicles (EVs) hold strong diagnostic potential, yet conventional isolation methods such as ultracentrifugation and size-exclusion chromatography (SEC) involve manual handling steps and show substantial run-to-run variability, hindering clinical translation. This study presents the Vesicle Disk (VDisk), a centrifugal microfluidics-based EV purification platform that combines cation-exchange chromatography, sequential filtration, and multimodal chromatography for automated, label-free EV isolation from up to 1 mL of plasma. VDisk configurations differing in filter membrane and plasma volume (0.1–1.0 mL) are benchmarked against SEC for yield, purity, reproducibility and robustness. VDisk matches SEC in EV yield and exceeds it in EV/contaminant ratios at reduced input volumes, while achieving greater reproducibility (intra-donor CV < 5% for CD9 and CD81, versus up to 16% for SEC) and maintaining consistent yield and contaminant removal under both fasting and postprandial sampling conditions. The platform is application-tunable: processing up to 1 mL of plasma maximizes EV yield and concentration, whereas processing 0.5 mL achieves approximately 2–3-fold higher EV/total protein ratios than SEC. These findings establish the VDisk as an automated, robust and adaptable alternative to existing EV isolation methods for both research and clinical translation.

## 1 Introduction

Extracellular vesicles (EVs) are lipid bilayer-enclosed nanoparticles that are released by all cell types, mediating intercellular communication and contributing to various physiological and pathological processes ^1^. EVs reflect the molecular state of their cells of origin by transporting proteins, nucleic acids, and other biomolecules to recipient cells, enabling their use in diagnostics, drug delivery, and regenerative medicine ^2–4^.

Among biofluids, blood-derived plasma and serum are particularly promising for EV-based diagnostics ^5^. Since most downstream analytical assays require the separation of EVs from other components, a reliable and efficient isolation method is essential to ensure accurate and reproducible results ^6^. The lack of standardized and automated EV purification technologies is a major hurdle that needs to be overcome to unlock the use of EVs in clinical applications ^7^. EV isolation exploits differences in size, density, charge, and molecular composition, and established methods rely on one or more of these to separate EVs from contaminants ^8^. Differential ultracentrifugation, density gradient centrifugation, and size-exclusion chromatography (SEC) are the most commonly used isolation methods ^1^. However, the presence of abundant lipoproteins and plasma proteins, which share similar size, charge or density with EVs, leads to co-isolation of these confounding particles, introducing biases in EV analyses and hindering the comparability of results ^9^. Lipoproteins are a particular challenge: they are present in blood at concentrations up to 10^6^-fold higher than EVs and overlap with EVs in size and other physical properties. Depleting these particles is one of the main challenges in EV purification ^10^.

Microfluidics-based EV isolation technologies offer advantages over conventional methods such as faster processing and automation potential ^11^. However, many current implementations still suffer from poor purity due to co-isolation of proteins and lipoproteins ^5^. Immunoaffinity-based microfluidic platforms provide improved selectivity but are limited to EV subpopulations expressing specific surface markers, an inherent limitation given the absence of universal EV markers. In general, most microfluidic approaches yield only small amounts of EVs, insufficient for many downstream applications ^12–14^.

Centrifugal microfluidics has emerged as a promising technology for process automation due to its inherent operational advantages ^15^. Centrifugal forces, generated in a rotating cartridge, drive fluids through sequential processing steps. Unlike pressure- or syringe-pump-driven systems ^16^, centrifugal microfluidic platforms do not require external tubing or actuators, reducing system complexity and lowering the risk of contamination or operational error ^17^. In addition, well-established unit operations are available—such as metering, valving and mixing—that facilitate automation of complex assay workflows within a compact and closed system architecture ^15,18^.

Several EV isolation methods have been developed using centrifugal microfluidics. For example, Exodisc-B and Exodisc-P rely solely on filtration to isolate EVs from blood or plasma; however, their size-based separation strategy results in limited purity ^19^. Alternatively, a platform that incorporates functionalized membranes (Exo-CMDS) can achieve higher purity by targeting specific EV subpopulations, although this typically comes at the cost of lower recovery ^20^. The SpinEx system represents one of the most technologically advanced approaches reported to date. The cartridge processes 150 µL of whole blood, capturing EVs and labeling them for 16 protein markers ^21^.

This study introduces the Vesicle Disk (VDisk), a centrifugal microfluidic cartridge co-developed with a multimodal EV isolation protocol. By integrating cation-exchange chromatography (CEX), sequential filtration (SeqF), and multimodal chromatography (MMC), the VDisk can purify EVs from up to 1 ml of plasma. It enables label-free EV isolation with high run-to-run reproducibility, offering a robust automated platform for clinical research and translational applications. Unlike the SpinEx system, the VDisk is not designed for a specific diagnostic application but as a standardizable, automated, label-free EV purification tool for clinically relevant plasma volumes. In this study we assess the VDisk’s recovery of GFP-EVs spiked into plasma, test different plasma input volumes to define the best compromise between EV yield, purity and purification speed, and stress-test the VDisk’s robustness and reproducibility against lipemic samples. We compare the VDisk’s performance against the widely used IZON qEV original SEC columns.

## 2 Results

### 2.1 VDisk’s multimodal EV purification workflow

The VDisk integrates a multimodal isolation workflow consisting of three sequential steps (Figure 1): 1. cation-exchange chromatography (CEX); 2. sequential filtration including EV concentration (SeqF); 3. multimodal chromatography (MMC).

**Figure 1.**
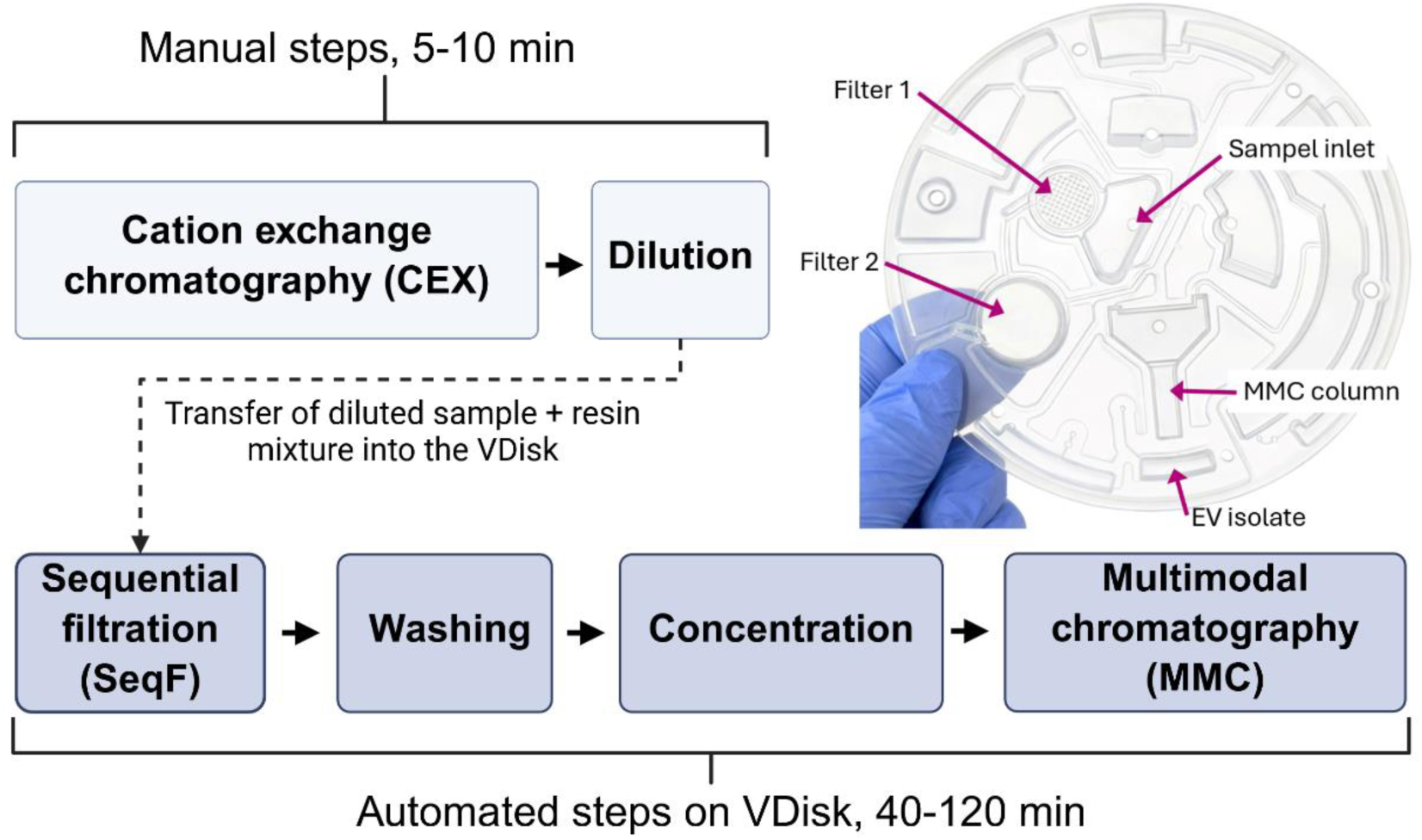
Overview of the VDisk workflow for EV isolation. The protocol integrates cation-exchange chromatography (CEX), sequential filtration, and multimodal chromatography (MMC) to eliminate contaminants based on charge and size. The CEX and dilution steps are performed manually and off-disk (5-10 min), while the subsequent steps are automatically executed on the VDisk (40–120 minutes, depending on sample volume). The top-right image shows an unprocessed VDisk.

In step 1, the plasma sample is incubated with CEX resin, which depletes positively charged plasma components, removing up to 85% of V/LDLs and 30% of soluble proteins within 5 min of incubation (Figure S2). This prevents membrane fouling in the next steps and allows up to 1 mL of plasma to be processed without clogging and with a significantly reduced filtration time. Subsequent dilution to a 3 mL final volume with PBS further improves the filtration and ensures fluidic operation of the VDisk.

In step 2, the mixture of diluted plasma and CEX resin is added to the VDisk mounted on the LabDisk-Player processing device and the automated processing is started (fluidic process: Figure S3, Table S1). During disk operation, the sample is filtered at low centrifugal force (max 140 × g). The first membrane—either a polycarbonate track-etched (PCTE) filter with 1.0 µm pores or a polyethersulfone (PES) filter with 0.22 µm pores—retains larger particles, including cell debris, protein aggregates, and the CEX resin. The resulting flow-through, containing EVs and smaller plasma components, passes to the second membrane (aluminum oxide, 0.02 µm pore size). This membrane retains EVs while smaller particles pass through. A subsequent wash with 2 mL of washing buffer removes most soluble proteins and HDLs. In step 3, the EV-enriched solution (∼700 µL) is then automatically transferred onto the integrated MMC column containing 500 µL of prepacked MMC resin (Figure S3). This final purification step eliminates residual proteins, nucleic acids, and other contaminants smaller than 400 kDa. Finally, purified EVs are pipetted out from a designated chamber.

### 2.2 GFP-EV Recovery

To assess EV recovery of the VDisk workflow in comparison with SEC under controlled conditions, GFP-EVs were spiked into healthy plasma from a single donor, purified in technical triplicate and quantified before and after purification. Biological variability is addressed separately in Sections 2.3 and 2.4 using plasma from three independent donors. Both VDisk configurations (VDisk_0.22µm and VDisk_1.0µm, see Section 5.1.2) were tested (Figure 2A). GFP-tagged EVs were spiked into plasma to a final concentration of 3 × 10¹⁰ particles/mL. For each VDisk configuration, 1 mL of spiked plasma was processed, while 0.5 mL of plasma was processed using SEC columns. Each VDisk run yielded approximately 700 µL of EV isolate and 4.3 mL of waste liquid. For SEC, the first three fractions (400 µL each, 1.2 mL total) were collected and pooled, since they contain 98% of total recovered EVs (Figure S4). EV recovery was calculated as the ratio of the absolute GFP-EV amount in the isolate to the absolute GFP-EV amount spiked into the initial sample.

**Figure 2.**
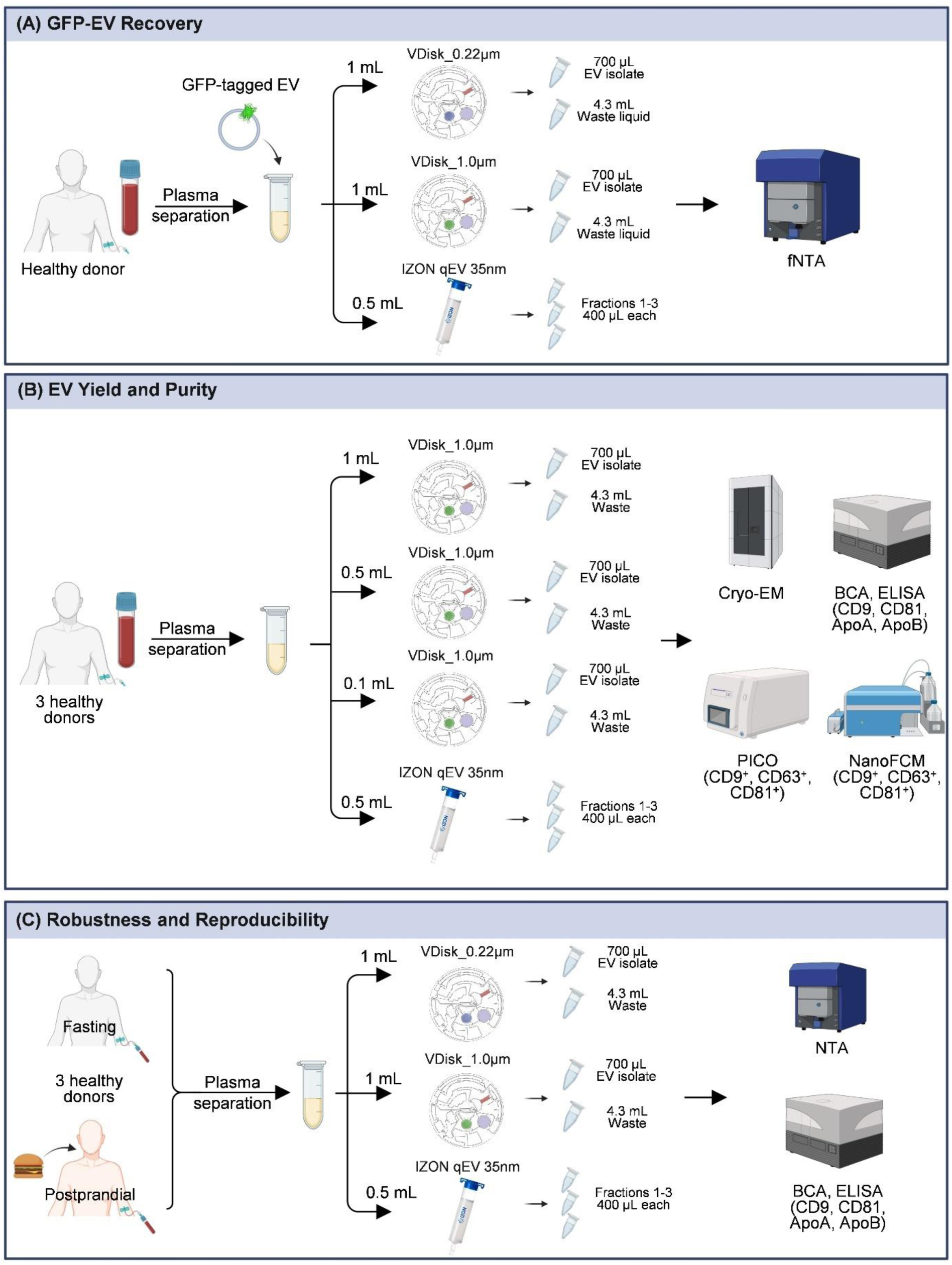
Overview of the experimental workflow. The study design comprises three major experiments: (**A**) GFP-EV recovery from spiked plasma, (**B**) EV yield and purity, analyzed with blood of three healthy donors and different disk loading volumes and, (**C**) benchmarking of robustness and reproducibility using fasting and postprandial plasma from the same three donors in three technical replicates. Different disk configurations and analysis methods were used as indicated. Created with BioRender.com.

As shown in Figure 3, VDisk_0.22µm achieved a recovery of 74.8% (±7.1%), while VDisk_1.0µm achieved a higher recovery at 84.3% (±2.2%). In comparison, the total recovery from SEC fractions 1–3 was 60.0% (±6.1%). Analysis of the VDisk waste liquid showed that less than 1% of GFP-EVs were present, indicating minimal EV loss through passage across the second filter membrane.

**Figure 3.**
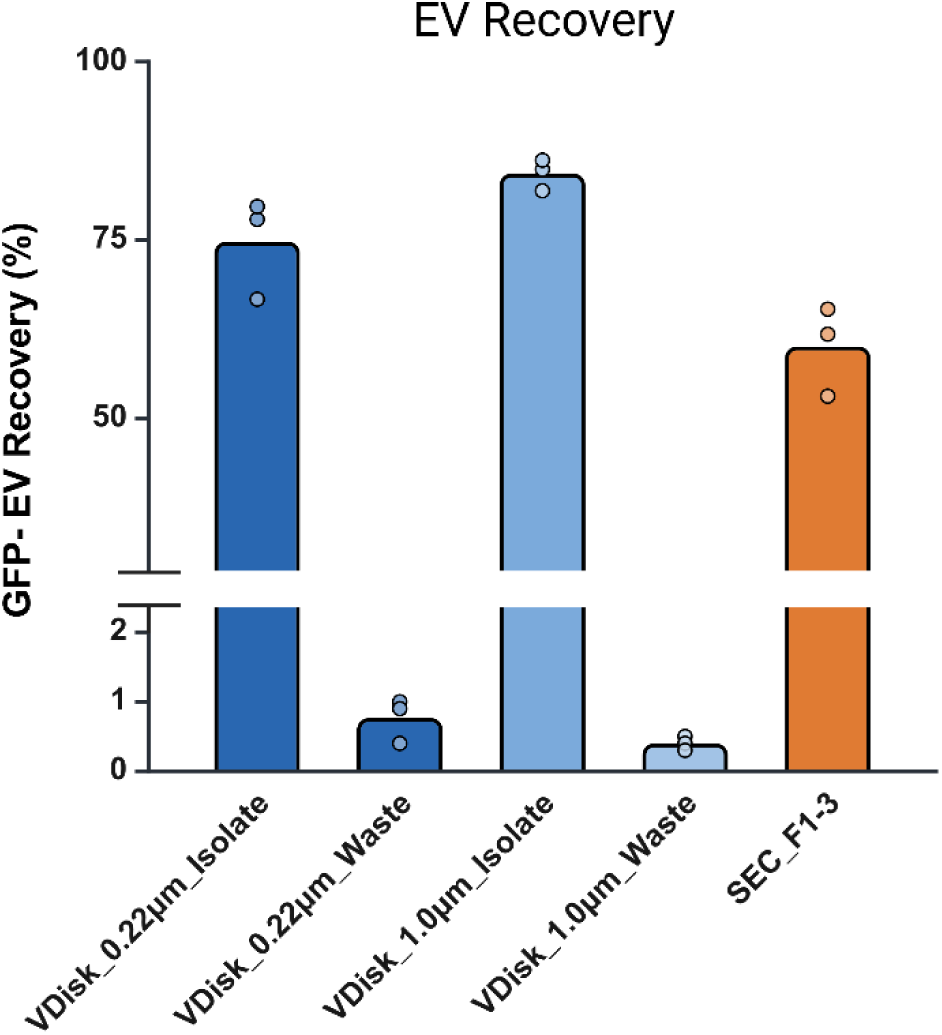
Recovery of GFP-tagged EVs from spiked plasma. EV recovery was quantified in the final isolates and corresponding waste fractions obtained using VDisk_0.22µm, VDisk_1.0µm, and SEC fractions 1-3. Plasma was obtained from one healthy donor with three independent technical replicates per condition; each data point represents a technical replicate. Created with BioRender.com.

### 2.3 EV Yield and Purity

To further assess the VDisk performance, yield and purity of endogenous EVs from healthy donor plasma were measured and related to contaminant levels using different methods (Figure 2B). As an additional parameter, the plasma volume loaded onto the VDisk was varied between 1.0 mL, 0.5 mL and 0.1 mL. Only VDisk_1.0µm, which uses the 1 µm pore size PCTE first membrane, was evaluated, since it had shown higher GFP-EV recovery. Table S2 outlines the corresponding experimental conditions.

EV isolates were analyzed using ELISA kits targeting CD9, CD81, ApoA, and ApoB, together with BCA protein assays, all of which have demonstrated high intrinsic reproducibility (Figure S5). To allow a fair comparison between different isolation methods—each starting with varying plasma volumes and resulting in different final isolate volumes—assay results were normalized, accounting for both the input and final volumes (see 5.2.10).

As shown in Figure 4A, VDisks loaded with 1.0 mL or 0.5 mL plasma resulted in comparable normalized CD81 yields, almost twice as high as SEC. In contrast, no significant difference between VDisk and SEC was observed when the disk was loaded with 0.1 mL plasma. Normalized yields also did not significantly differ between VDisks and SEC when measuring CD9 (Figure S6A).

**Figure 4.**
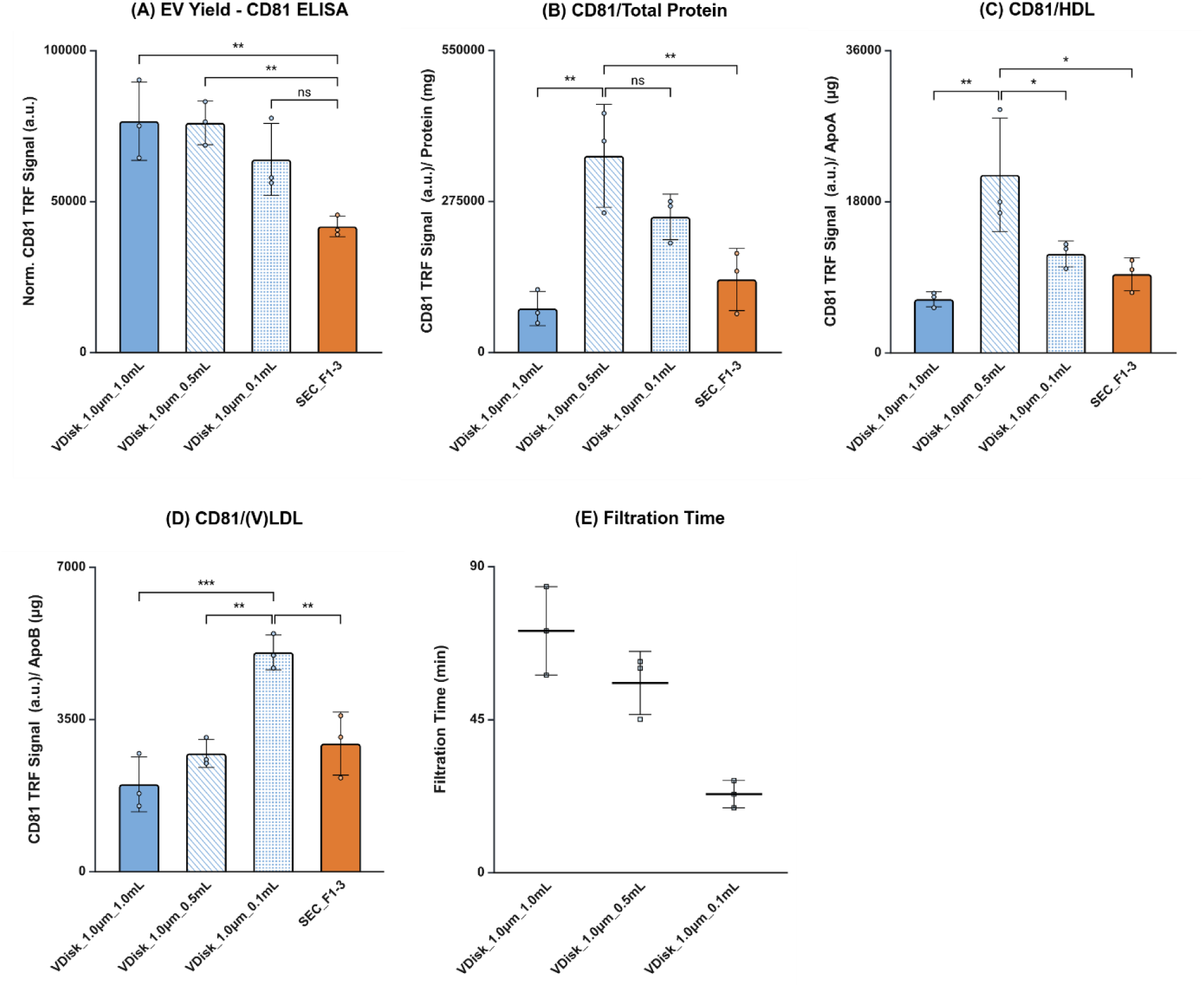
VDisk_1.0µm performance at varying plasma volumes compared with SEC. (**A**) EV yield (normalized CD81 TRF signal), (**B**) CD81 TRF signal per total protein, (**C**) CD81 TRF signal per ApoA (HDL marker), (**D**) CD81 TRF signal per ApoB (V/LDL marker), and (**E**) total filtration time (filter 1 plus filter 2). Bars show means from three healthy donors (biological replicates); each point represents an individual donor. Error bars indicate standard deviations. Statistical analysis: one-way ANOVA with Tukey’s multiple comparisons test (n = 3). Significance levels: ns = not significant, *p < 0.05, **p < 0.01, ***p < 0.001. Created with BioRender.com.

Figure 4B–D present EV purity as the ratio of CD81 signal to protein, HDL-associated ApoA, and V/LDL-associated ApoB contaminations, respectively. VDisks processing 0.5 mL of plasma achieved 2–3-fold higher purity than SEC for EV-to-total protein and EV-to-HDL ratios. When processing 0.1 mL of plasma, VDisk outperformed SEC in the EV-to-V/LDL ratio, with nearly a 1.5-fold improvement (Figure 4D). Figure S6 presents the same analysis using CD9 instead of CD81, showing consistent trends with slight differences in ratios.

Filtration time in the VDisk increased with plasma input volume (Figure 4E). For instance, 1 mL input volume required approximately 80 min to complete the filtration steps, whereas 0.5 mL and 0.1 mL required around 55 min and 30 min, respectively.

Figure 5 presents the normalized concentrations of EV subpopulations (CD9^+^, CD63^+^, and CD81^+^) measured using two single-EV analysis methods: NanoFCM, a flow cytometry platform optimized for small particles ^22^, and PICO, a novel digital immunoassay designed to quantify individual particles independently of size or contaminants ^23,24^. Both NanoFCM and PICO measurements indicate similar yields across different VDisk configurations and SEC for CD9, CD63, and CD81 containing EVs. Figure 5C shows the percentage of marker-positive nanoparticles in the isolates obtained by VDisk and SEC, as determined by NanoFCM. For CD9⁺ and CD81⁺ EVs, the percentage of marker-positive particles was higher for the VDisk loaded with 0.1 mL of plasma than for any other variant. However, the number of acquired events for this sample was low and fell below the acquisition target during the defined acquisition time, which was set equal for all samples to ensure comparability (Figure S12).

**Figure 5.**
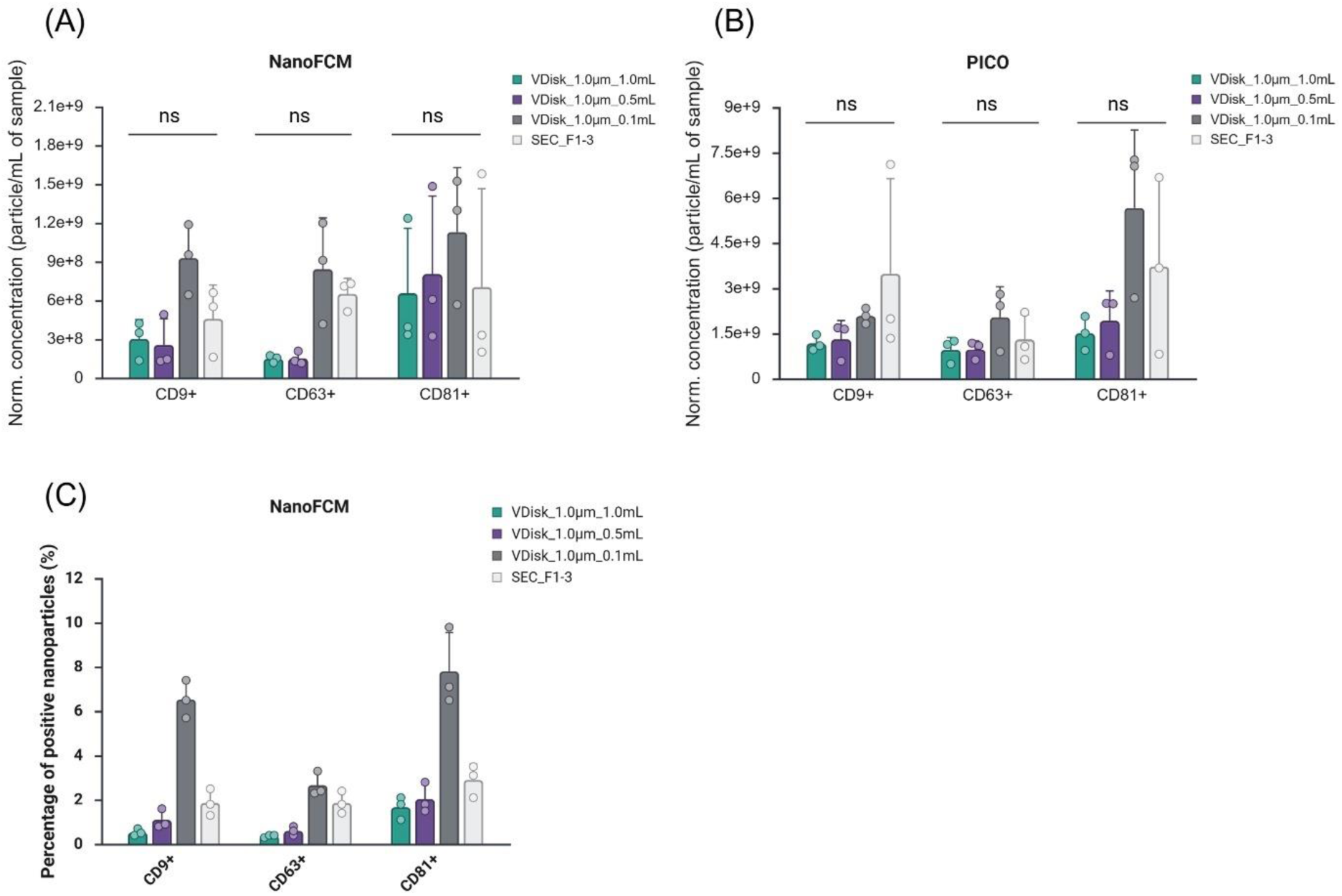
Single-EV analysis of subpopulation concentrations across different isolation methods. Normalized concentration of CD9-, CD63-, and CD81-positive EVs was quantified by (**A**) nanoFCM and (**B**) PICO. (**C**) Percentage of positive nanoparticles quantified by nanoFCM. Bars show means from three healthy donors (biological replicates); each point represents an individual donor. Error bars indicate standard deviations. Statistical analysis in A and B: two-way ANOVA with Tukey’s multiple comparisons test (n = 3). Significance levels: ns = not significant, *p < 0.05, **p < 0.01, ***p < 0.001. No statistical comparison was performed in due to low number of acquired events (C).

Cryo-EM images (Figure 6) confirm the presence of intact EVs in both VDisk and SEC isolates.

**Figure 6.**
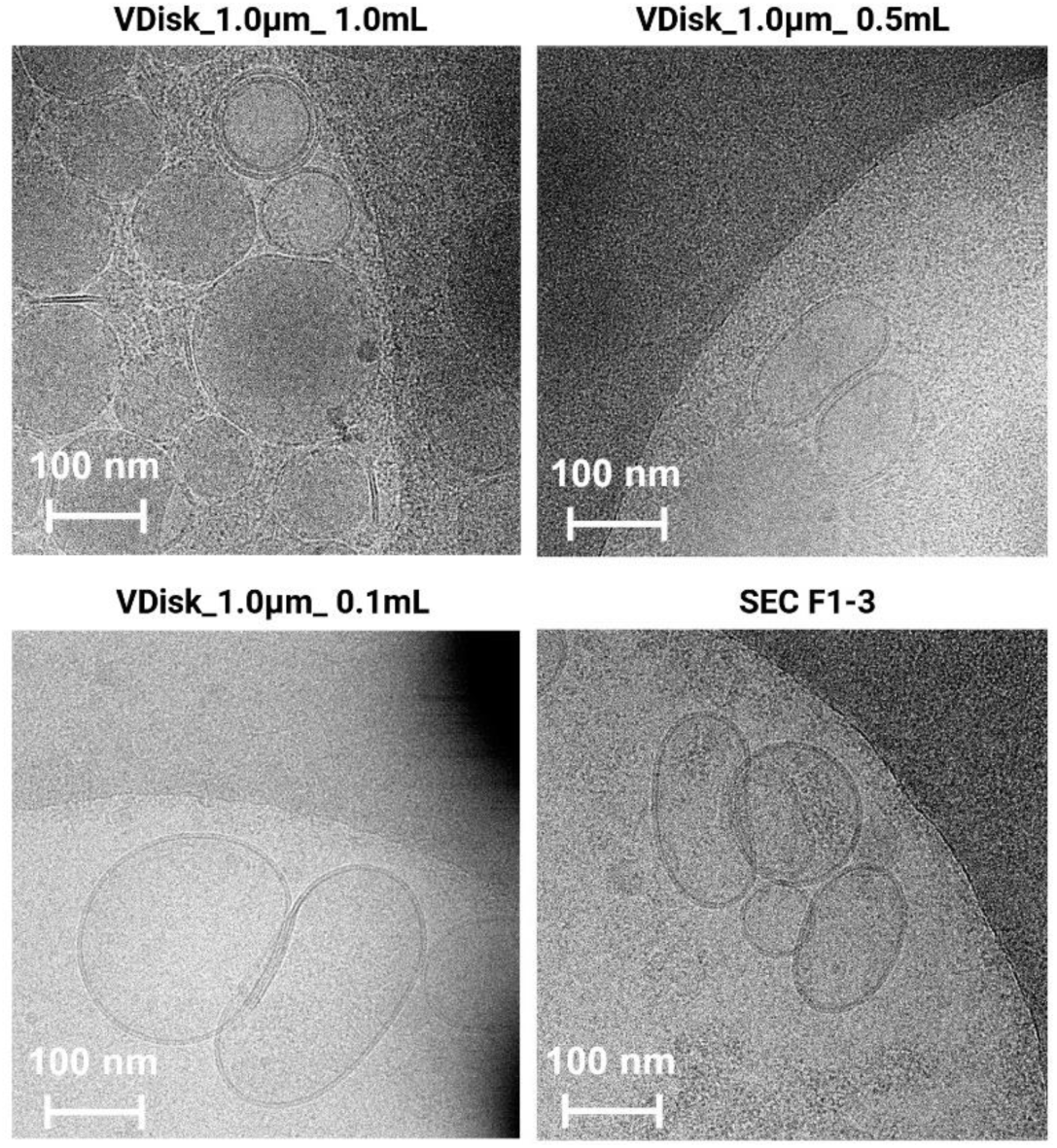
Cryo-EM images of EVs obtained using VDisk_1.0µm using different input volumes and SEC, all displaying intact vesicular morphology.

### 2.4 Robustness and Reproducibility

To further characterize the isolation performance and stress-test the robustness of the VDisk, blood samples were collected from three healthy donors in both fasting and postprandial states (4 hours after food intake, Figure 2C) on the same day. As shown in Figure S7A-B, postprandial plasma samples displayed higher optical density and a milky appearance, indicating increased lipid levels ^25,26^. Total particle concentration, CD9 and CD81 signal intensities, and protein and lipoprotein concentrations were assessed before isolation for all samples and are presented in Figure S7C-F. Performance of both VDisk_0.22µm and VDisk_1.0µm was evaluated loading 1 ml of these plasma samples in three technical replicates each (three separate VDisks per sample) and benchmarked against SEC.

Based on CD9 and CD81 ELISA results, both VDisk configurations produced yields with no significant differences observed between fasting versus postprandial plasma (Figure S8). A slight shift in populations was nevertheless apparent, with increased CD9 and decreased CD81 signals in postprandial plasma for all three donors.

Table 1 summarizes the mean intra-donor CV% values. Both VDisk configurations exhibited mean CVs below 5% for CD9 and CD81 in both fasting and postprandial states, showing minimal variation between the two conditions. On the other hand, SEC CV% values were notably higher (up to 16 %) and especially increased in postprandial samples (raw data presented in Figure S9). Similar results were obtained for total protein, ApoA and ApoB results, where VDisk was around twice as reproducible as SEC.

**Table 1.**
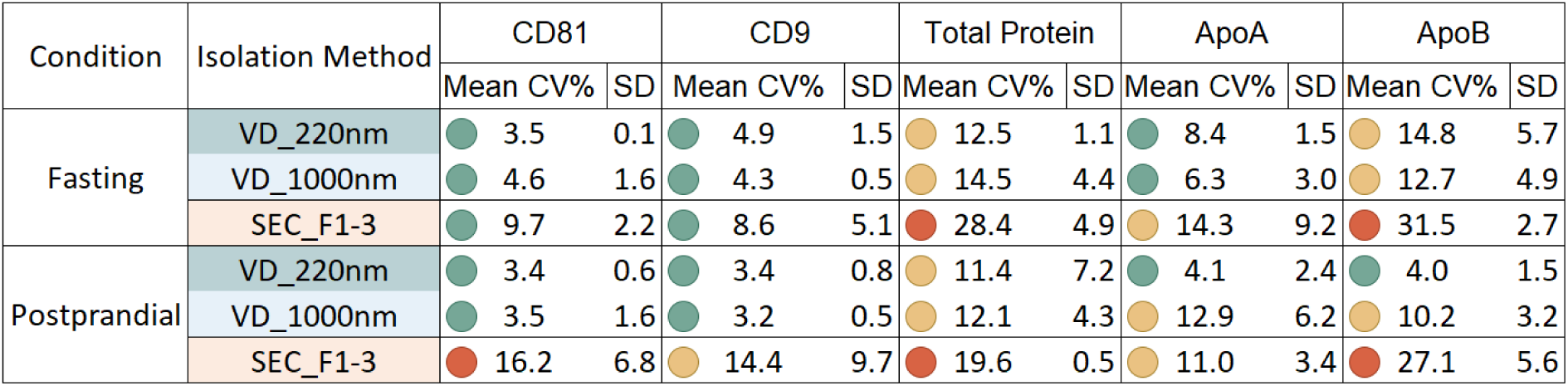
Reproducibility of EV yield and purity across isolation methods. Mean intra-donor coefficients of variation (CV, %) and corresponding standard deviations (SD) are shown for CD81, CD9, total protein, ApoA, and ApoB measurements under fasting and postprandial conditions (n = 3 healthy donors, each with three technical replicates). Color coding: green = CV < 10%, yellow = 10–15%, red ≥ 15%. Raw data shown in Figure S9.

| Condition | Isolation Method | CD81 |  | CD9 |  | Total Protein |  | ApoA |  | ApoB |  |
| --- | --- | --- | --- | --- | --- | --- | --- | --- | --- | --- | --- |
|  |  | Mean CV% | SD | Mean CV% | SD | Mean CV% | SD | Mean CV% | SD | Mean CV% | SD |
| Fasting | VD_220nm | ● 3.5 | 0.1 | ● 4.9 | 1.5 | ● 12.5 | 1.1 | ● 8.4 | 1.5 | ● 14.8 | 5.7 |
|  | VD_1000nm | ● 4.6 | 1.6 | ● 4.3 | 0.5 | ● 14.5 | 4.4 | ● 6.3 | 3.0 | ● 12.7 | 4.9 |
|  | SEC_F1-3 | ● 9.7 | 2.2 | ● 8.6 | 5.1 | ● 28.4 | 4.9 | ● 14.3 | 9.2 | ● 31.5 | 2.7 |
| Postprandial | VD_220nm | ● 3.4 | 0.6 | ● 3.4 | 0.8 | ● 11.4 | 7.2 | ● 4.1 | 2.4 | ● 4.0 | 1.5 |
|  | VD_1000nm | ● 3.5 | 1.6 | ● 3.2 | 0.5 | ● 12.1 | 4.3 | ● 12.9 | 6.2 | ● 10.2 | 3.2 |
|  | SEC_F1-3 | ● 16.2 | 6.8 | ● 14.4 | 9.7 | ● 19.6 | 0.5 | ● 11.0 | 3.4 | ● 27.1 | 5.6 |

Figure 7A–D illustrates the removal efficiency of protein, HDL, V/LDL and total particles using VDisk_0.22µm, VDisk_1.0µm, and SEC under both fasting and postprandial conditions. All methods demonstrated high removal efficiency, with no statistically significant differences between fasting and postprandial conditions. Both VDisk configurations achieved ≥98% total protein removal, ≥99% HDL-associated ApoA removal, ≥93% V/LDL-associated ApoB removal, and ≥92% total particle removal. IZON SEC performed similarly, with slightly better performance in V/LDL and total particle removal. The size distribution determined by NTA (focus on 80–150 nm particles) remained nearly unchanged between the fasting and postprandial isolates (Figure 7E-F). The NTA data shown are not normalized to sample or elution volumes and instead report the final concentrations of VDisk preparations and pooled SEC fractions. Particle concentrations are substantially higher in VDisk preparations than in SEC elution fractions. Three factors contribute: the VDisk was loaded with twice the sample volume (1.0 mL versus 0.5 mL), SEC dilutes the sample during elution, and the VDisk concentrates EVs on the second filter membrane.

**Figure 7.**
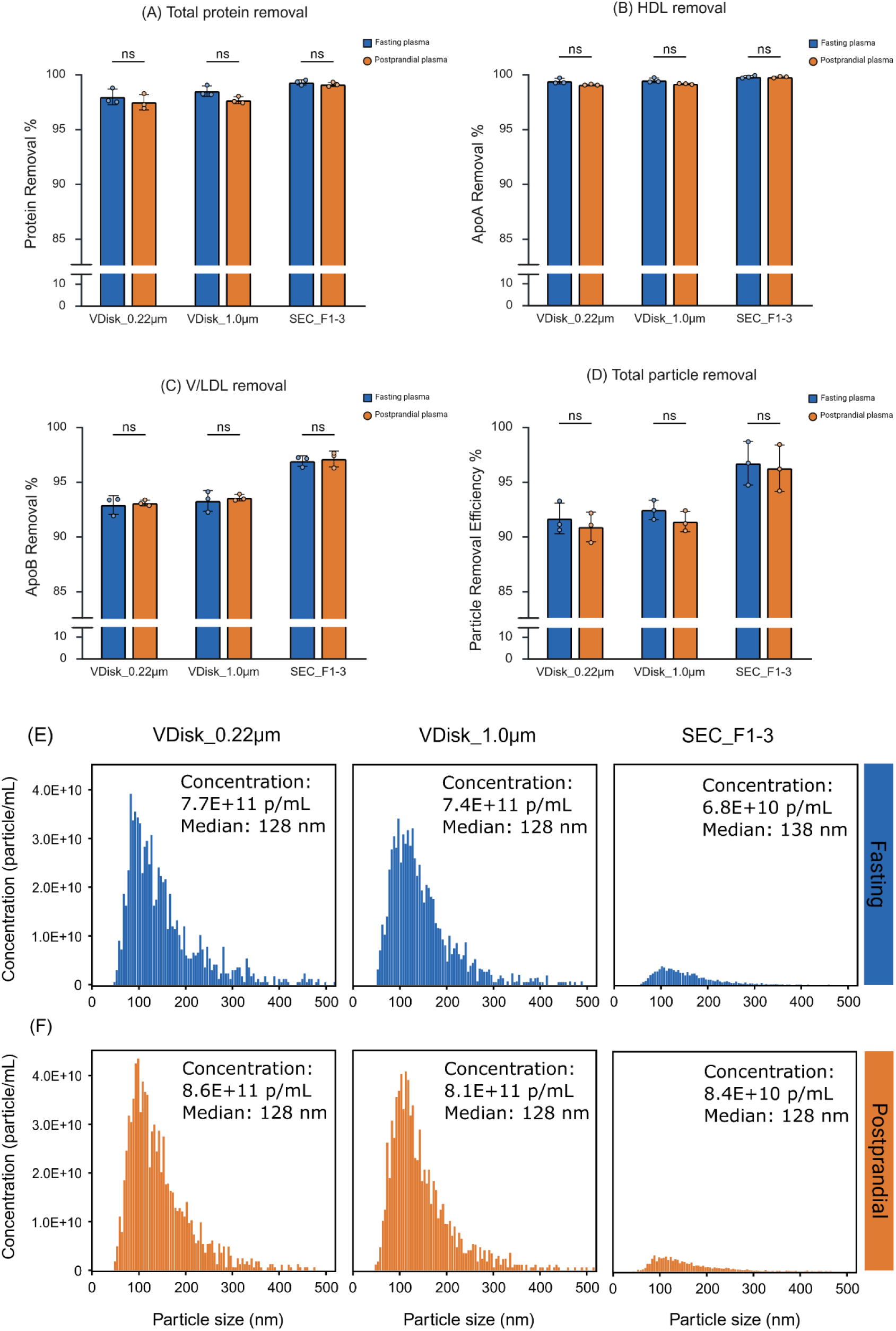
Contaminant removal and size distribution of EV isolates across methods. (**A**) Total protein removal measured by the BCA assay, (**B**) ApoA removal (HDL marker) measured by ELISA, (**C**) ApoB removal (V/LDL marker) measured by ELISA, and (**D**) total particle removal measured by NTA; bars represent means from three healthy donors (biological replicates), each data point reflects the average of three technical replicates, and error bars indicate standard deviations. Statistical analysis: two-way ANOVA with Tukey’s multiple comparisons test (n = 3). Significance levels: ns = not significant, *p < 0.05, **p < 0.01, ***p < 0.001. (**E**, **F**) NTA particle size distribution profiles from fasting and postprandial plasma, respectively. Created with BioRender.com.

Variation in filtration time among technical replicates was minimal, with CVs below 10% for each method, indicating the operational consistency of the VDisk platform (Figure S8C–E). Notably, VDisk_0.22µm maintained stable filtration times between fasting and postprandial samples, indicating robustness to varying plasma conditions. In contrast, VDisk_1.0µm showed an approximately 25% increase in filtration time when processing postprandial plasma.

## 3 Discussion

In this study we introduced the VDisk for automated EV extraction after a single manual pre-incubation step, integrating cation-exchange chromatography (CEX), sequential filtration (SeqF) and multimodal chromatography (MMC) for EV purification from plasma and compared it to IZON qEV original 35 nm columns. The VDisk was tested with two different first-membrane materials and loaded with plasma of different volumes and lipid content.

In the recovery experiments with GFP-EV spiked plasma (section 2.2), VDisks with PES 0.22 µm or PCTE 1.0 µm membranes both achieved higher EV recovery than SEC, with VDisk_1.0µm (PCTE) reaching 84.3% compared to 60.0% for SEC (Figure 3). VDisk_1.0µm slightly outperformed VDisk_0.22µm (PES), likely because of structural differences between the membranes used for the first filtration step. The PES 0.22 µm membrane has a smaller cut-off, greater thickness (∼150 µm) and an interconnected pore architecture (Figure S10A), which may retain larger EVs. In contrast, the PCTE 1.0 µm membrane has larger pores, reduced thickness (∼20 µm), and a uniform pore size (Figure S10B), facilitating more efficient EV passage. Notably, less than 1% of GFP-EVs were detected in the waste liquid, confirming minimal loss through the secondary 0.02 µm aluminum oxide membrane, used on both VDisk configurations for concentrating and washing EVs. The remaining 15–25% EV loss is likely attributable to nonspecific adsorption to the Disk’s polymer and filter or MMC resin particles, whose surface morphology is shown in Figure S10C. Because EV adsorption losses are expected to increase as competing protein is depleted, a surface-blocking component was added to the Disk’s washing buffer.

In general, it should be noted that recovery values obtained using spiked-in GFP-EVs are only partly translatable to endogenous EVs. Because their surface composition is altered and they have already been purified once from cell culture supernatant (centrifugation, syringe filtration, TFF; see Section 5.1.7), they do not reproduce the heterogeneity of endogenous plasma EVs.

Next, we investigated the yield of endogenous EVs in healthy donor plasma and evaluated how VDisk plasma input volumes (1.0 ml, 0.5 ml, 0.1 ml) affect the balance between EV yield and purity. Normalized VDisk yield was almost twice as high as SEC in the CD81 ELISA when 1.0 mL or 0.5 mL plasma was loaded onto the Disk (Figure 4A). However, differences were non-significant for CD9 EVs (Figure S6A). The lower normalized yield achieved with 0.1 mL plasma is likely caused by greater EV loss through adsorption. Additionally, the relative performance of VDisk and SEC appears to depend on the tetraspanin marker used for quantification. One possible explanation is that EV populations differ in their expression of tetraspanin subtypes, and that different isolation methods preferentially recover distinct EV subtypes depending on their underlying isolation principles.

EV yields obtained with each isolation method were additionally assessed using two single-EV analysis techniques, NanoFCM and PICO. As shown in Figure 5A–B, no statistically significant differences were observed between VDisk and SEC for any of the CD9, CD63, or CD81 markers. However, the high variability of these assays and their inherent biases when analyzing samples with differing purity might mask differences between isolation methods.

Notably, EV yields reported by PICO (Figure 5B) were consistently higher than those measured by NanoFCM (Figure 5A). This discrepancy is likely attributable to the size detection limit of NanoFCM, which is constrained to 40–150 nm particle diameters, whereas PICO is not subject to this restriction. The PICO assay shows strong potential as a tool for EV detection, especially since it is capable of identifying both single-marker–positive and dual-marker–positive populations (Figure S11). Future studies will be needed to systematically investigate these methodological differences and their implications for cross-platform comparability.

When assessing EV/contaminant ratios, the VDisk was superior to SEC in most comparisons. VDisks processing 0.5 mL plasma showed a 2–3-fold higher purity ratio compared to SEC for EVs/total protein and EVs/HDL (Figure 4B–C and S6B–C). When focusing on EV/(V)LDL ratios, the VDisks processing 0.1 mL plasma performed best, achieving approximately 1.5-fold higher purity than SEC (Figure 4D and S6D). VDisks processing 1 mL plasma yielded the least pure isolates across all conditions but provided the highest EV concentration due to the larger processed sample volume, making it advantageous for downstream applications where higher EV numbers are required and moderate protein or lipoprotein contamination is less critical.

In addition to ELISA, purity was assessed by NanoFCM as the percentage of CD9-, CD63- and CD81-positive particles relative to the total particle population (Figure 5C). Across the 1.0 mL and 0.5 mL conditions, which met the target acquisition range, marker-positive fractions remained below 2%. This low percentage reflects the overwhelming abundance of lipoproteins, which are present at several orders of magnitude higher concentrations than EVs in plasma and is in line with previous publications ^10^. Consequently, even after substantial lipoprotein depletion they still numerically exceed EVs.

For the 0.1 mL and SEC isolates, event counts fell below the range required for reliable enumeration (Figure S12), and we therefore do not draw conclusions from their positivity values, even though the 0.1 mL condition showed the numerically highest percentage. The control panel — blank (dilution buffer), unstained, antibody-only and isotype controls — confirmed signal specificity and excluded relevant background (Figure S13).

To better contextualize these relative purity metrics, we additionally present the absolute contaminant load in the final EV isolates (Figure S14). Notably, purity does not decrease linearly across the tested input sample volumes. Processing 1 mL of plasma resulted in the highest level of contamination, significantly higher than for the 0.5 mL and 0.1 mL input volumes. Increasing the initial sample volume proportionally increases the absolute number of particles entering the system. Larger particles, such as V/LDL that are not removed during the CEX step, can accumulate on the 0.02 µm membrane, forming a filter-cake that reduces filtration efficiency and limits contaminant removal during the washing steps and following MMC, ultimately leading to reduced final EV purity.

Finally, to stress-test the VDisk, we compared loading 1 mL of fasting or postprandial plasma. Physiologically, chylomicrons are the dominant lipoproteins released into circulation a few hours after food intake ^27^. As shown in Table 1, even for these chylomicron-rich samples the EV yield (CD9 and CD81 ELISA) was highly reproducible when the same sample was purified three times for both VDisk_0.22µm and VDisk_1.0µm, with CV values consistently below 5%. This reproducibility exceeded that of SEC, which showed CVs up to 16%. SEC reproducibility especially suffered in postprandial samples, whereas the VDisk was not negatively affected. This leads to the conclusion, that SEC is stronger influenced by sample quality compared to the VDisk.

Notably, SEC was operated with the manufacturer’s automatic fraction collector, so this comparison is against a partially automated reference method rather than fully manual SEC. Overall, EV yields were in a similar range in fasting and postprandial samples (Figure S8A–B). EV populations appear slightly shifted in postprandial samples, however, trending towards an increase in CD9 and a decrease in CD81.

Beyond EV isolation yield, postprandial plasma had no significant impact on the removal of total protein, HDL, V/LDL, and total particles (Figure 7A–D). In addition, the particle size distribution of VDisk isolates remained unchanged, and the median particle size measured by NTA shifted by only approximately 10 nm in SEC isolates. This was unexpected, as a pronounced increase in particle numbers had been anticipated for both VDisk isolates, particularly VDisk_1.0µm, and SEC under postprandial conditions. Chylomicrons span a size range of 0.08–1.2 µm ^28^, which would allow a substantial fraction to pass through the PCTE 1.0 µm membrane used in VDisk_1.0µm, while being largely retained by the PES 0.22 µm membrane in VDisk_0.22µm. This difference in first membrane cut-off size likely explains why filtration times for VDisk_0.22µm remained consistent between fasting and postprandial samples, whereas filtration times for VDisk_1.0µm increased by approximately 25% under postprandial conditions (Figure S8C–E). Particles passing through the 1.0 µm first membrane presumably accumulate on the downstream 0.02 µm aluminum oxide membrane, which dominates total filtration time.

The absence of a detectable increase in particle numbers in postprandial VDisk_1.0µm isolates can be attributed to a combination of effective lipoprotein removal and analytical limitations of particle counting. The CEX step of the VDisk purification workflow efficiently depletes positively charged lipoproteins (Figure 4D, S2A), indicating that the majority of postprandial lipoproteins are removed prior to downstream analysis. While classical SEC only purifies based on size, IZON claims that their improved Gen 2 resin is also able to retain positively charged particles. This may explain why IZON SEC columns perform well in removing V/LDLs, which fall in the same size range as EVs ^29^.

A further factor is that residual large lipoproteins such as chylomicrons are quantified with substantially lower precision by NTA analysis, since the applied SOP100 protocol is validated for EV-sized particles (80–150 nm). Differences in large lipoprotein abundance between fasting and postprandial samples may therefore fall within the measurement uncertainty of the total particle counts rather than being resolved.

## 4 Conclusion

In summary, the VDisk automates plasma EV isolation after a single 5-minute manual pre-incubation, delivering intra-donor CVs below 5% for CD9 and CD81 where IZON SEC reached 16%, while matching SEC in yield and surpassing it in contaminant ratios when comparable input volumes (0.5 ml) are used. This combination — high reproducibility, automation and purity — is what positions the VDisk for standardized EV biomarker studies.

By leveraging centrifugal microfluidics in combination with a multimodal label-free isolation workflow, the VDisk performs comparably to IZON SEC in EV yield and achieves higher EV-to-contaminant ratios at reduced input volumes, while SEC retains a small advantage in absolute V/LDL and total particle removal. Its compatibility with variable sample volumes, tolerance to physiological heterogeneity (e.g., postprandial lipemia), and low CV values underscore its translational potential for both research and clinical contexts. The VDisk also delivers EVs at higher concentrations than SEC, which often requires an additional concentration step. The Disk yields a final isolate volume of 700 µL with CD9⁺, CD63⁺ or CD81⁺ EV concentrations in the range of 10⁸–10^10^ EVs/mL, depending on the starting sample volume and donor variability (Figure 5A–B). Further concentration, down to 500 µL, is possible at the cost of longer filtration times. In practical terms, a single VDisk isolate contains sufficient EV material to support multiple downstream assays. In this study, a single VDisk isolate was used to perform BCA, several ELISAs, NanoFCM, fNTA, PICO and cryo-EM analyses.

As with any integrated EV purification method, the VDisk platform is subject to certain limitations. First, plasma throughput is restricted to 1 mL and only one sample can be processed at a time, limiting sample throughput. In addition, the workflow requires a manual incubation step with the CEX resin, introducing 5 minutes hands-on work to the isolation protocol. Finally, EV losses due to adsorption or nonspecific interactions with the resin surfaces cannot be fully prevented. Nonetheless, these limitations are outweighed by the advantages of high EV/contaminant ratios and high reproducibility.

Several emerging methods aim to automate plasma EV purification ^30,31^, but many still suffer from substantial variability in the final isolates, often attributable to plasma pre-processing steps ^32^. A future iteration of the VDisk will address this limitation by integrating plasma separation and CEX on the cartridge itself, enabling a fully automated workflow starting from fresh whole blood. Because plasma preparation is itself time-consuming and operator-dependent, moving it onto the cartridge is where the largest reduction in hands-on time and user influence is expected. In parallel, the development of a dedicated LabDisk-Player equipped with a built-in fluorescence detector will enable on-disk sample and isolate characterization, such as quality control assessments (e.g., hemolysis or lipemia), as well as the detection of EV-associated markers like protease activity ^33^, thereby supporting sample-to-answer EV analyses. Together, these developments would position the VDisk to translate laboratory workflows into clinical practice and, ultimately, to support EV-based point-of-care diagnostics.

## 5 Materials and Methods

### 5.1 VDisk Based EV Isolation

#### 5.1.1 VDisk Isolation Protocol

An overview of the VDisk isolation workflow is presented in Figure 1. First, 1 mL of plasma sample was mixed at a 1:1 ratio with pre-washed CEX resin (Section 5.1.4) in a Protein LoBind® 5.0 mL screw cap tube (Eppendorf, Cat# 0030122356). The mixture was vortexed and incubated for 5 minutes at room temperature (RT) on a rotary mixer (RS-RD 5, Phoenix Instrument) set to 1500 rpm. Afterwards, 2 mL of phosphate-buffered saline (PBS, pH 7.4; Gibco, Thermo Fisher Scientific, Cat# 20012027) was added to dilute the sample. The resulting slurry—comprising plasma, resin, and diluent—was vortexed and subsequently loaded into the inlet chamber of the VDisk.

Details of the VDisk’s automated fluidic operation are summarized in Table S1. The protocol was executed using a customized LabDisk-Player (Euler Player, 2nd Generation; BioFluidix GmbH, Germany) (Figure S1A–B). Briefly, filtration was performed at a constant rotational frequency of 25 Hz (maximum RCF: 140 × g), followed by the addition of 2 mL washing buffer (WB, section 5.1.5), with intermittent shake-mode mixing (25/5 Hz, acceleration rate of 15 Hz/s) to prevent membrane clogging. Filtration proceeded until approximately 700 µL of EV-enriched solution was collected on the second membrane. This fraction was then transferred to the integrated MMC column, which was pre-packed with 500 µL of pre-washed MMC resin (Section 5.1.4). The final isolate was collected from the designated chamber as illustrated in Figure 1, by puncturing the sealing foil.

#### 5.1.2 VDisk Configurations

VDisk was evaluated in two main configurations, which differed in the membrane material and pore size used for the first filtration step. The VDisk_0.22µm configuration employs a 0.22 µm polyethersulfone (PES) membrane filter (Sartorius, Cat# 15427-EP), while the VDisk_1.0µm configuration uses a 1.0 µm polycarbonate (PCTE) membrane filter (Sterlitech, Cat# 1270092). In both configurations, the second membrane is a 0.02 µm aluminum oxide Anodisc™ 25 filter with support ring (Whatman, Cat# 6809-6002).

#### 5.1.3 VDisk Preparation

VDisk was designed using SolidWorks 2018 software (Dassault Systèmes) and fabricated via injection molding (Kaiser Ingenieurbüro GmbH, Germany) from a cyclic olefin copolymer (COC) blend containing COC 6013 (TOPAS, Cat# 6013S-04) and COC 8007 (TOPAS, Cat# 8007S-04) with a ratio of 70% to 30%, respectively. Membranes were thermally bonded to the cartridge using an in-house built sealing unit (Figure S1C), and the open side of the disk was closed using a sealing press (Wickert hydraulic presses) with COC foil (120 µm thick, Tekniplex) as shown in Figure S1D.

#### 5.1.4 Chromatography Resins Preparation

Fractogel® EMD SO₃⁻ (M), a strong cation-exchange (CEX) resin (Millipore, Cat# 1168820010), was washed three times with PBS prior to use and stored at 4 °C for up to two weeks. The same washing procedure was applied to Capto™ Core 400, a multimodal chromatography (MMC) resin (Cytiva, Cat# 17372402).

#### 5.1.5 Washing Buffer (WB) Preparation

PBS supplemented with 0.5% (v/v) SmartBlock™ (Candor, Cat# 113125) and 0.005% (v/v) Poloxamer 188 (Sigma-Aldrich, Cat# P5556-100ML) was used as washing buffer (WB).

#### 5.1.6 Size Exclusion Chromatography (SEC) Based EV Isolation

SEC was used as a reference method for EV isolation, employing qEV original 35 nm columns (IZON), designed for 500 µl sample volume, with Gen2 resin together with the Automatic Fraction Collector (AFC; IZON). The protocol was performed according to the manufacturer’s instructions with minor modifications. Briefly, columns were equilibrated with 50 mL of buffer prior to loading 500 µL of plasma. Subsequently, 3–5 fractions of 400 µL each were collected, with the void volume defined at 2.9 mL.

#### 5.1.7 Preparation of Fluorescently Labeled EVs

For the EV recovery experiments, GFP-tagged EVs were isolated from HT1080-CD9-GFP cells, following the protocol described by ^27^. Briefly, cells were cultured in T-175 cm² flasks at 37 °C with 5% CO₂ and 95% humidity in antibiotic-free Roswell Park Memorial Institute (RPMI) 1640 medium (Gibco/Thermo Fisher Scientific), supplemented with 10% fetal bovine serum (FBS; Thermo Fisher Scientific) and 20 mM L-glutamine (Sigma-Aldrich). Approximately 200 million cells were grown to 80% confluency, washed twice with Dulbecco’s phosphate-buffered saline (1× DPBS; Gibco/Thermo Fisher Scientific), and then incubated in serum-free RPMI medium for 24 hours. The conditioned medium was collected and sequentially centrifuged at 800 × *g* for 5 minutes and 5000 × *g* for 25 minutes to remove cells, debris, and apoptotic bodies. The resulting supernatant was passed through a 0.22 µm PES syringe filter (Millex-GP, Merck KGaA) and concentrated to ∼1 mL using tangential flow filtration (TFF-easy; HansaBiomed Life Sciences Ltd). The concentration of GFP-tagged EVs was quantified by fluorescent nanoparticle tracking (fNTA) prior to spiking into plasma samples.

#### 5.1.8 Blood Collection and Plasma Preparation

Whole blood was collected from three healthy male donors (written informed consent has been obtained) using S-Monovette® Citrate 9NCtubes (Sarstedt, Germany), and processed using a two-step centrifugation protocol ^34^ to prepare plasma. Briefly, 10 mL of whole blood was collected and gently inverted five times to ensure thorough mixing with the anticoagulant. Within 30 minutes of collection, samples were centrifuged at 2,500 × g for 15 min at room temperature. The resulting plasma layer was carefully transferred to a 15 mL Falcon tube, leaving approximately 1 cm plasma above the buffy coat/plasma interface. The plasma was then subjected to a second centrifugation step at 2,500 × g for 15 min at room temperature. After centrifugation, the clarified plasma was transferred to a fresh 15 mL Falcon tube, again leaving the lower ∼1 cm of plasma untouched to avoid contamination from cellular debris. Centrifugation was performed using a Heraeus Megafuge 16R (Thermo Scientific) equipped with a swing-out bucket rotor. Immediately after plasma preparation, the samples were aliquoted into volumes suitable for downstream experiments and stored at −80 °C. Prior to freezing, turbidity was measured as a pre-analytical quality control step to evaluate changes in postprandial plasma lipid level (Section 5.2.1).

### 5.2 Analytical Methods

#### 5.2.1 Assessment of Lipid Level

To evaluate the lipid level, all samples were diluted 10-fold with PBS. The absorbance of each diluted sample was measured at 610 nm using a microplate reader (TECAN) in 96-well microliter plates (Greiner non-binding).

#### 5.2.2 Nanoparticle Tracking Analysis (NTA)

For quantification of particle concentration and size distribution, QUATT Nanoparticle Tracking Analysis (NTA; Particle Metrix GmbH, Germany) was performed using Zeta Navigator software (version 1.4.2.1). EVs were measured on the day of purification and stored on ice. Freeze-thaw cycles were avoided. Samples were diluted to achieve 80–200 particles per frame for valid acquisition. Scatter mode settings were: sensitivity 80, shutter 90, trace length 12 s, frame rate 30 fps, 7 positions, minimum size 10 nm, maximum size 1000 nm, laser 488 nm, and temperature 25 °C. Fluorescence mode settings were: sensitivity 95, shutter 50, trace length 12 s, frame rate 30 fps, 7 positions, minimum size 10 nm, maximum size 1000 nm, laser 488 nm, filter 500 nm, and temperature 25 °C. Zeta potential measurements were conducted using the following settings: sensitivity 80, shutter 90, cycle frames 30, cycle amount 4, frame rate 30 fps, 2 positions (SL1 and SL2), dielectric constant 78.434, Henry factor 1.5, laser 488 nm, and temperature 25 °C. All parameters (size distribution, particle concentration, and zeta potential) were measured using 1:10 diluted PBS in Milli-Q H₂O (Stakpure system, OmniaTap XS basic). Final size distribution analysis was performed with the Phonups software ^35^. Measurements followed SOP100, a standard operating procedure calibrated against 100 nm reference particles, whose validated range is 80–150 nm. Within this range, particle size and concentration are determined with high precision. Particles outside the range are still detected and are included in the reported size distributions and total particle counts, but their size and concentration carry correspondingly larger uncertainty. No size gating was applied.

#### 5.2.3 EV Surface Marker Quantification

To compare the EV isolation yield across different isolation methods, CD9 and CD81 ELISA kits (Cell Guidance Systems, Cat# EX501 and EX503) were used following the manufacturer’s protocol. Briefly, streptavidin-coated 96-well plates were incubated with biotinylated anti-CD9 or anti-CD81 antibodies (2 ng/µL, 100 µL/well, 1 h, RT). After washing, 100 µL of samples or PBS (blank) were added and incubated (1 h, RT). Prior to loading, all samples were diluted to ensure that analyte concentrations fell within the linear dynamic range of each assay (Figure S15). Plates were washed and incubated with europium-labeled anti-CD9 or anti-CD81 antibodies (0.25 ng/µL, 100 µL/well, 1 h, RT). Following additional washes, europium fluorescence intensifier (EFI, 100 µL/well) was added and incubated for 15 min. Fluorescence was measured using a time-resolved fluorescence (TRF) microplate reader (excitation 340 nm, emission 615 nm).

#### 5.2.4 Total Protein Quantification

Protein concentrations were determined using the Pierce™ BCA Protein Assay Kit (Thermo Fisher Scientific, Cat# 23225 and 23227) following the manufacturer’s protocol without modifications.

#### 5.2.5 Lipoprotein Quantification

HDL concentrations were determined using an ApoA1 ELISA kit (R&D Systems, Cat# DAPA10) and V/LDL concentrations using an ApoB ELISA kit (R&D Systems, Cat# DAPB00). In both cases, the manufacturer’s protocol was followed without modifications.

#### 5.2.6 Cryo-EM

Cryo-EM grids were prepared by applying 3 µL of sample onto glow-discharged holey carbon grids (Quantifoil R 1.2/1.3, Cu 300 mesh; 25 s at 10 mA, Pelco EasiGlow). Grids were blotted using a Vitrobot Mark IV (22 °C, 100% humidity, 6 s, blot force 5) and plunge frozen in liquid ethane. When higher particle density was required, multiple rounds of sample application and blotting were performed prior to freezing. Data collection was performed on a 300 kV Titan Krios G4 (Thermo Fisher Scientific) equipped with a Selectris energy filter and Falcon 4i direct electron detector. Images were acquired at 105,000× nominal magnification with a total dose of 40 e⁻/Å² and a defocus setting of 3 µm.

#### 5.2.7 Scanning Electron Microscopy (SEM)

Filter membranes or dried MMC resins were placed on aluminum stubs using carbon adhesive tape and sputter-coated with a thin gold–palladium layer. SEM imaging was performed on a Tescan MAIA3 XMH electron microscope equipped with a secondary electron detector, operated at an accelerating voltage of 2 kV.

#### 5.2.8 Nano-Flow Cytometry

Nanoflow cytometry was performed using a Nano Analyzer (NanoFCM Co., Ltd., Nottingham, UK) equipped with 488 nm and 640 nm lasers. Prior to sample measurements, the instrument was calibrated with 250 nm silica calibration beads (QS3003, NanoFCM Co.) at a concentration of 2.11 × 10¹⁰ particles/mL, which also served as the reference for particle concentration determination. Size calibration was carried out using monodisperse silica beads (S16M-Exo, NanoFCM Co. Ltd.) of four different sizes (68 nm, 91 nm, 113 nm, and 155 nm).

For immunofluorescence staining, VDisk 1mL, VDisk 0.5mL, VDisk 0.1mL and SEC samples were labeled for specific surface markers using the following antibodies: anti-human CD9 (H19a)-FITC (BioLegend, 312108), anti-human CD63 (H5C6)-FITC (353007, BioLegend), and anti-human CD81 (5A6)-APC (BioLegend, 349509). Samples were incubated with antibodies for 1 hour at room temperature.

Data acquisition was performed over a 1-minute period at a constant sample pressure of 1.0 kPa. For analysis, all samples were diluted in particle-free 1× PBS to achieve a target count of 2,500–12,000 events per minute. Particle concentration and size distribution were determined using NF Profession v2.08 software (NanoFCM Co. Ltd.).

To ensure signal specificity, the following controls were included in each experiment: unstained samples, blank control, antibody-only controls (CD9, CD81, CD63), and isotype controls. Mouse IgG1-FITC, κ and Mouse IgG1κ-APC were used as isotype controls for CD9, CD63 and CD81 respectively. Following staining, all samples and controls were diluted 1:200 in PBS immediately prior to acquisition.

#### 5.2.9 Protein Interaction Coupling (PICO) Assay

The detection of surface markers in intact EVs was performed according to the manufacturer’s instructions. In brief, labeled antibodies against the tetraspanins CD9, CD63 and CD81 (#PICO-001011 to #PICO-001013, PICO BioScience) were mixed overnight at 4 °C with the EV sample at a working concentration of 4 × 10^-11^ M. In every experiment, an antibody binding control (ABC), which contains the antibody mix but lacks the sample of interest, was included.

Before the dPCR readout, the samples were diluted to aim for an average antibody concentration per partition of 0.15, as recommended by ^23^. The dPCR was performed using QIAGEN’s QIAcuity Digital PCR System according to PICO BioScience’s manual, using the PICO probes matching the DNA labels conjugated to the antibodies (PICO BL, P8, N6 and O7 probes; #PICO-000070 to 73, PICO BioScience).

The raw dPCR results were analyzed with the PIQUANT software (PICO BioScience) as described in ^23,24^. The results include a correction for the ABC values (which account for offsets in the dPCR data caused by signal dropouts or incorrect clustering), as well as a normalization against the lambda values, to compensate for handling errors. At least three technical replicates were conducted for each sample.

#### 5.2.10 Statistical Analysis

To compare methods differing in plasma input and final isolate volume, all assay values were blank-subtracted and normalized to both volumes according to Equation (1):

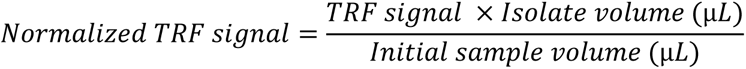

The same normalization was applied to the BCA, ApoA, ApoB, nanoFCM and PICO data. Data are presented as mean ± standard deviation. Throughout, n refers to the number of biological replicates (independent healthy donors); each donor sample was processed in three technical replicates, i.e. three separate VDisks or SEC columns. No data points were excluded and no outlier test was applied. Intra-donor coefficients of variation (CV) were calculated per donor across technical replicates, and Table 1 reports the mean of the three donor CVs. Comparisons across plasma input volumes and SEC (Figures 4, S6) were analyzed by one-way ANOVA; comparisons involving two factors (Figures 5, 7, S7, S8, S11) were analyzed by two-way ANOVA. The ANOVA F-test is omnibus and non-directional; pairwise comparisons were two-sided and corrected by Tukey’s multiple comparisons test, with α = 0.05. Significance is indicated as ns = not significant, *p < 0.05, **p < 0.01, ***p < 0.001. Given n = 3 per group, normality was assumed rather than formally tested. One-way and two-way ANOVA with Tukey multiple comparisons test were performed using BioRender Graphing (BioRender, Toronto, ON, Canada; https://www.biorender.com; RRID:SCR_018361), powered by R version 4.2.2.

## 6 Acknowledgments

This work was supported by the German Federal Ministry of Research, Technology and Space (BMFTR) project *KI-VesD-2* (Grant No. 03LW0339). We gratefully acknowledge support of the Facility for Extracellular Vesicle Analysis and Liquid Biopsy (EV-Core), Medical Center – University of Freiburg (registered with the German Research Foundation [DFG] under RI_00612), and the financial support it received from the Faculty of Medicine – University of Freiburg (project 2025/A13-Naz). The Electron microscopy data were collected at the Microscopy and Image Analysis Platform (MIAP), University of Freiburg: Cryo-EM Facility of the University of Freiburg (RRID: SCR_025860) supported by the German Research Foundation [DFG], project no. 506518771), and Martin Helmstaedter, Transmission Electron Microscopy Core Facility of the Medical Center University of Freiburg.

The authors acknowledge the use of Claude Opus 5 (Anthropic, August 2026) for assistance in grammar and language editing. All experimental design, data generation, analysis, and interpretation were performed solely by the authors.

## 7 Conflict of Interest and Disclosure

The VDisk platform is not a commercial product. At this stage, it remains an advanced laboratory prototype under active development and validation.

At present, a subset of the co-authors (Ehsan Mahmodi Arjmand, Gustav Grether, Jan Lüddecke, and Nils Paust) are evaluating the possibility of establishing a future startup to commercialize the VDisk platform. This initiative remains exploratory, and no company has been formed nor has any commercial activity begun.

Pablo Sánchez-Martín is employed as Head of R&D at PICO Bioscience and contributed to the study by performing the PICO assays.

## 8 Data Availability Statement

The data that support the findings of this study are available from the corresponding author upon reasonable request.

