## Supporting Information for "VDisk: A Microfluidic Cartridge for Automated and Reproducible Label-Free Isolation of Extracellular Vesicles from Plasma"

### Tables

***Table S1. VDisk fluidic protocol.***

| Step | Procedure |
|---|---|
| 1 | Pipette 3 mL of sample mixture into the VDisk. |
| 2 | Add the washed Capto Core 400 resin into the MMC column. |
| 3 | The VDisk rotated at 25 Hz with an acceleration of 15 Hz/s until the liquid level on the second membrane reaches the middle of the chamber (maximum RCF: 140 × g). |
| 4 | 10 cycles of shake-mode mixing between 25 Hz and 5 Hz with an acceleration of 15 Hz/s is performed. |
| | Add 1 mL of washing buffer into the sample inlet. |
| 5 | The VDisk is rotated at 25 Hz with an acceleration of 15 Hz/s until the liquid level on the second membrane reaches the middle of the chamber (maximum RCF: 140 × g). |
| 6 | 10 cycles of shake-mode mixing between 25 Hz and 5 Hz with an acceleration of 15 Hz/s is performed. |
| 7 | Add 1 mL of washing buffer into the sample inlet. |
| 8 | The VDisk rotated at 25 Hz with an acceleration of 15 Hz/s until the liquid level on the second membrane reaches the middle of the chamber (maximum RCF: 140 × g). |
| 9 | 10 cycles of shake-mode mixing between 25 Hz and 5 Hz with an acceleration of 15 Hz/s is performed. |
| 10 | The VDisk is rotated at 25 Hz for 5 seconds, followed by 45 seconds at 5 Hz. |
| 11 | Repeat steps 9–10 for a total of 15 cycles. During this process, the EV solution is transferred to the MMC column and incubated with Capto Core 400 resin for 15 minutes. |
| 12 | The VDisk is rotated at 30 Hz for 10 seconds to transfer the remaining EV solution from the MMC column into the EV collection chamber. |
| 13 | The VDisk is stopped, the EV isolate can be pipetted out from the EV collection chamber. |

***Table S2. Summary of VDisk_1.0µm variants processing different plasma input volumes.*** *The table lists the plasma input, resin, diluent volume, and the final output volume for each VDisk_1.0µm evaluated in this study. CEX resin volumes refer to settled resin and are not counted in the 3.0 mL liquid volume loaded into the VDisk (initial sample + diluent)*

| VDisk configuration | Initial sample volume | CEX resin volume | Diluent volume | First membrane cut-off Size | Second membrane cut-off Size | MMC resin volume | Final EV isolate volume |
|---|---|---|---|---|---|---|---|
| VDisk_1.0µm_1.0mL | 1.0 mL | 1.0 mL | 2 mL | 1.0 µm | 0.02 µm | 0.5 mL | 0.7 mL |
| VDisk_1.0µm_0.5mL | 0.5 mL | 0.5 mL | 2.5 mL | 1.0 µm | 0.02 µm | 0.5 mL | 0.7 mL |
| VDisk_1.0µm_0.1mL | 0.1 mL | 0.1 mL | 2.9 mL | 1.0 µm | 0.02 µm | 0.5 mL | 0.7 mL |

### Figures

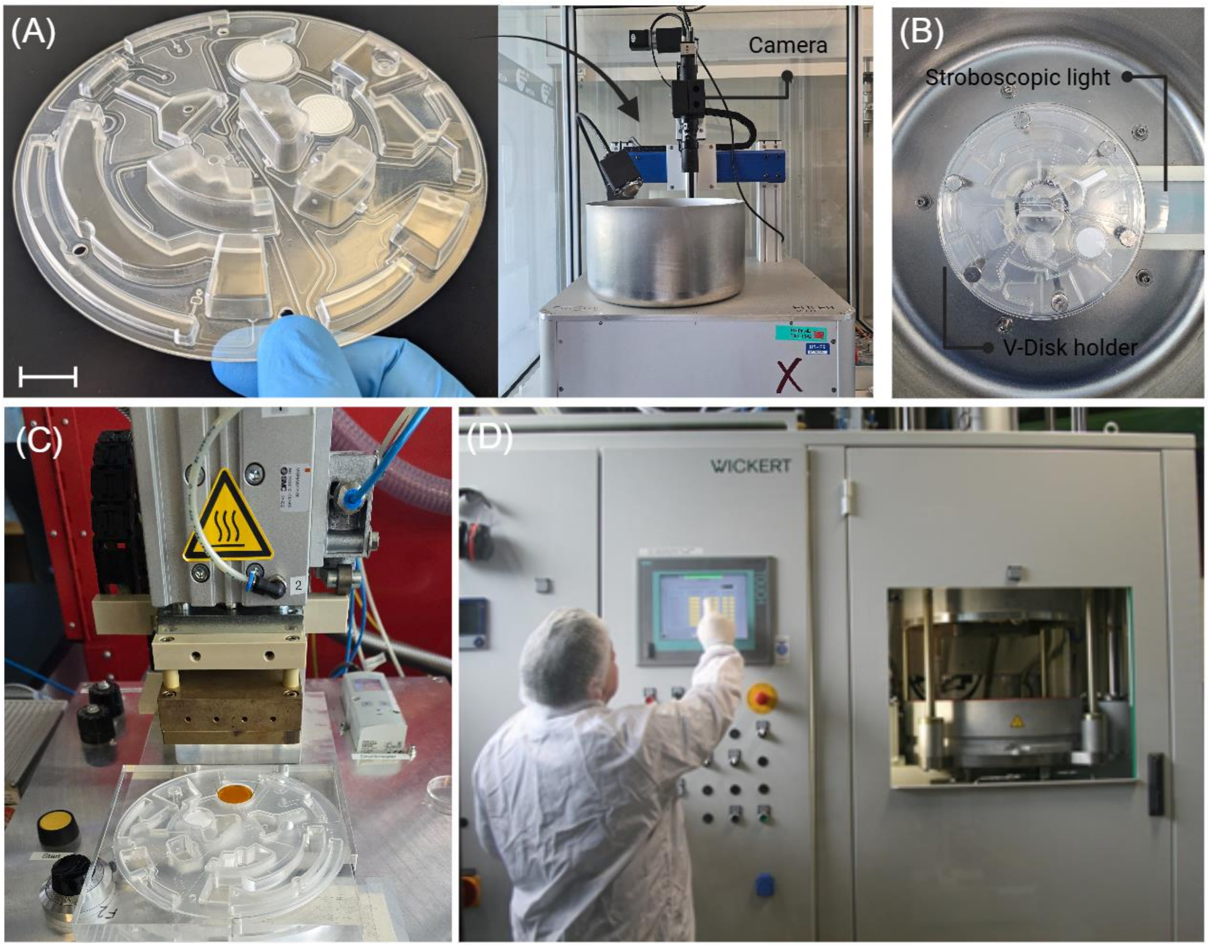


***Figure S1.*** *(A) VDisk and LabDisk-Player equipped with a camera for visualization of the fluidic operations during rotation. (B) VDisk mounted on the LabDisk-Player ready for operation. (C) Custom-built filter integration unit (sealing parameters: 180 °C, 4.5 bar, 8 s for 0.02 µm and 1.0 µm pore size membranes; 18 s for 0.22 µm pore size membrane). (D) VDisk sealing device (Wickert; 128 °C, 10 kN, 30 s) used to seal the VDisk with a COC foil.*

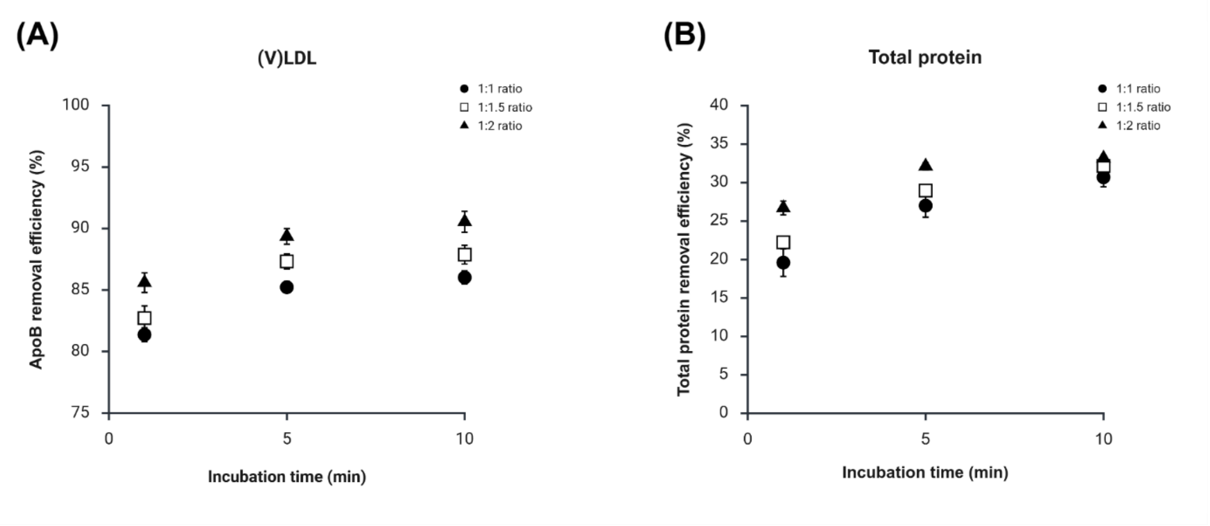


**Figure S2.** Impact of incubation time and sample-to-resin ratio on contaminant removal in the cation-exchange chromatography (CEX) step. (A) ApoB removal (marker for V/LDL) measured by ELISA, and (B) total protein removal measured by the BCA assay. Each data point represents the mean of a technical triplicate, error bars indicate standard deviations.

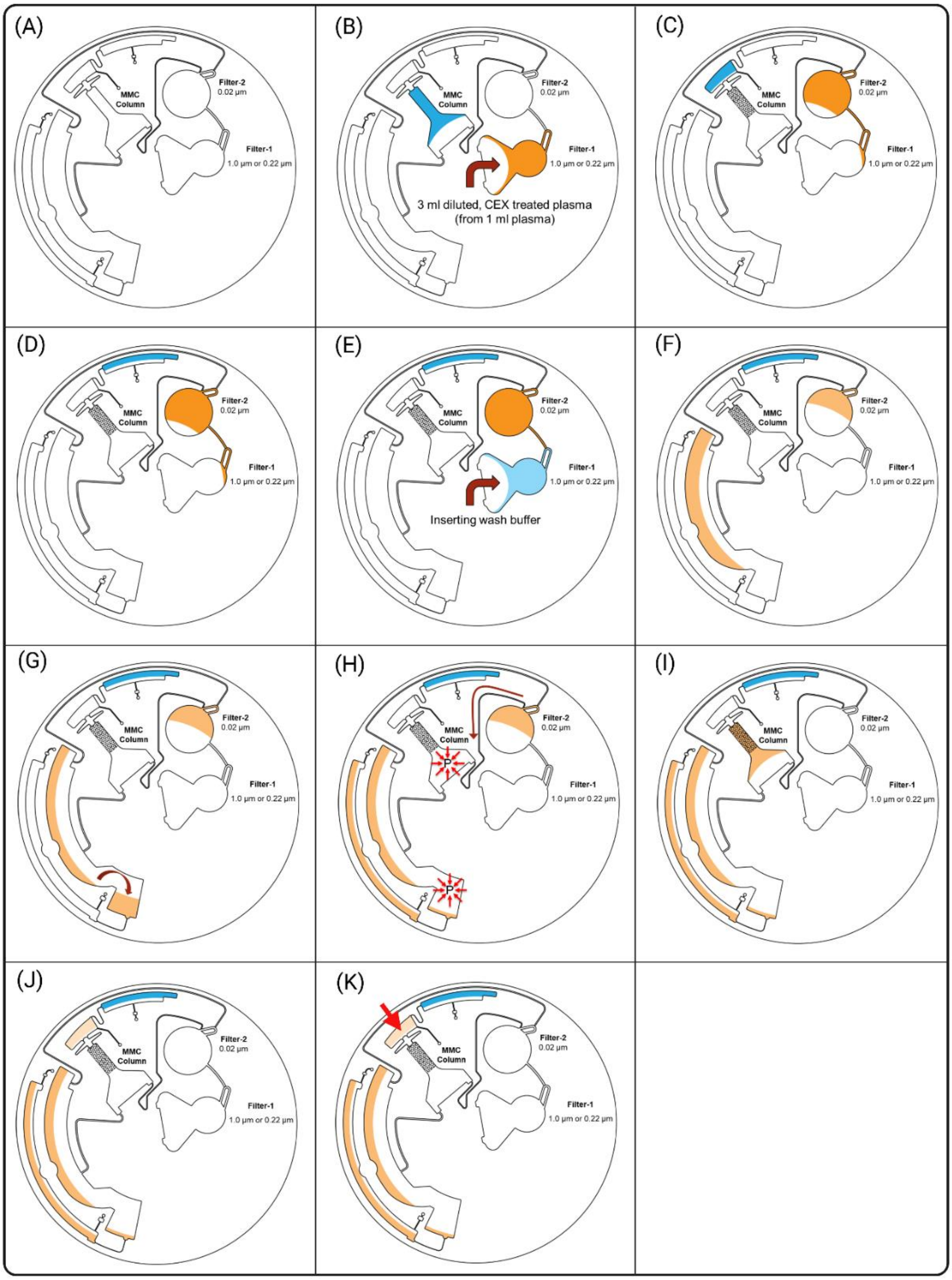


***Figure S3. Illustration of VDisk fluidic operation.*** *(A) VDisk layout. (B) Loading of the plasma sample and MMC resin into the VDisk. (C–D) Centrifugation at 25 Hz for 2 minutes packs the MMC resin into the column while approximately 650 µL PBS (blue) is transferred to the EV collection chamber and then to the waste chamber; during this step, the sample passes through filter 1. Spinning continues until the liquid level on filter 2 reaches mid-level. (E–F) Addition of 2 mL washing buffer followed by continued filtration with intermittent shake-mode mixing until the liquid level again reaches mid-level on filter 2. (G–H) Waste liquid is swapped to the opposite side of the waste chamber, and underpressure is generated by pumping it into the adjacent chamber. (I–K) EVs are pumped from filter 2 into the MMC column, incubated with the MMC resin, and the purified EVs are collected in the chamber indicated by red arrows.*

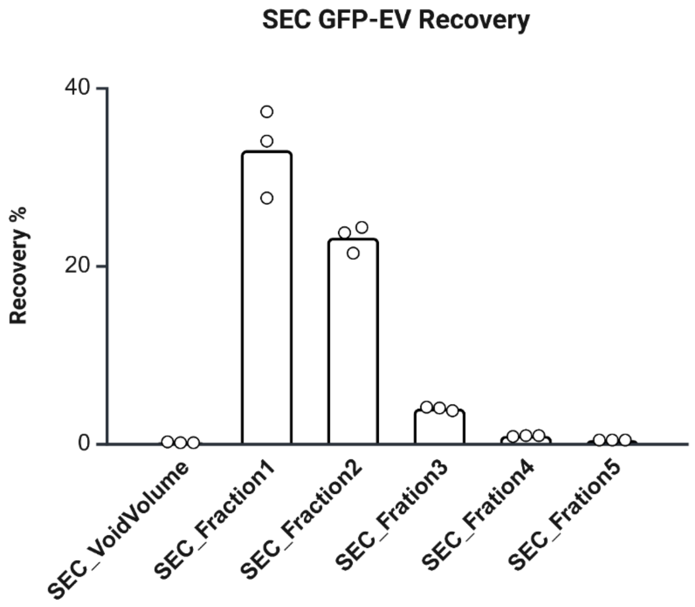


***Figure S4. Recovery profile of GFP-EVs across SEC fractions.*** *GFP-EV recovery (%) was quantified by fNTA in the void volume (2.9 mL) and five sequential SEC fractions (400 µL each). Bars represent mean values, with individual data points indicating technical replicates. Created with BioRender.com.*

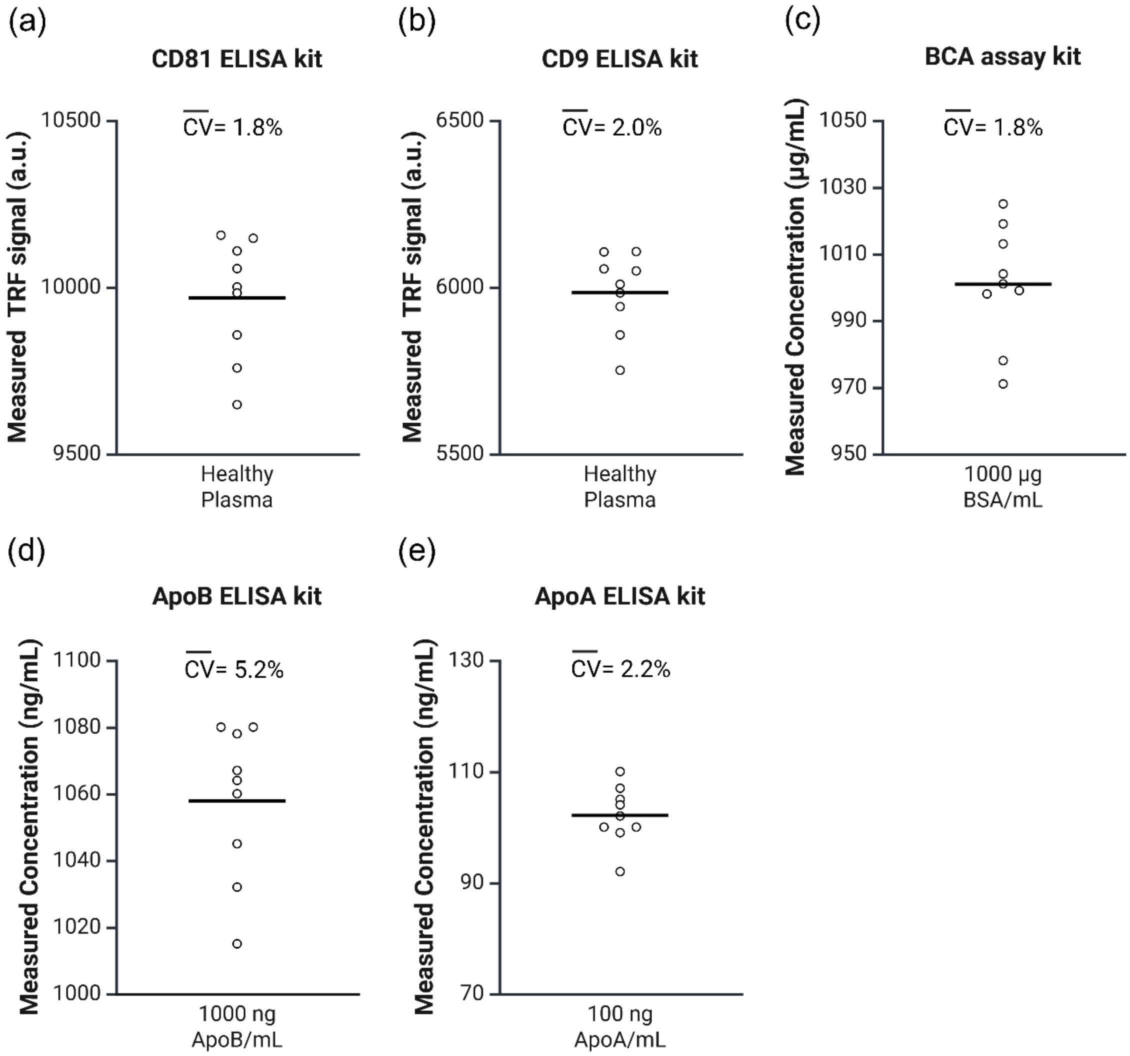


***Figure S5. Reproducibility of analytical assays used for EV and contaminant quantification.*** *Each assay was tested using nine technical replicates prepared at the same concentration to assess intra-assay variability. The CD81 and CD9 ELISA kits were evaluated with healthy plasma, the BCA assay with 1000 µg/mL BSA, and the ApoA and ApoB ELISA kits with 100 ng/mL ApoA and 1000 ng/mL ApoB standards, respectively. The narrow distribution of measured values across replicates indicates high analytical precision. Mean coefficients of variation* ($\overline{CV}$) *were calculated to benchmark assay reproducibility. Created with BioRender.com.*

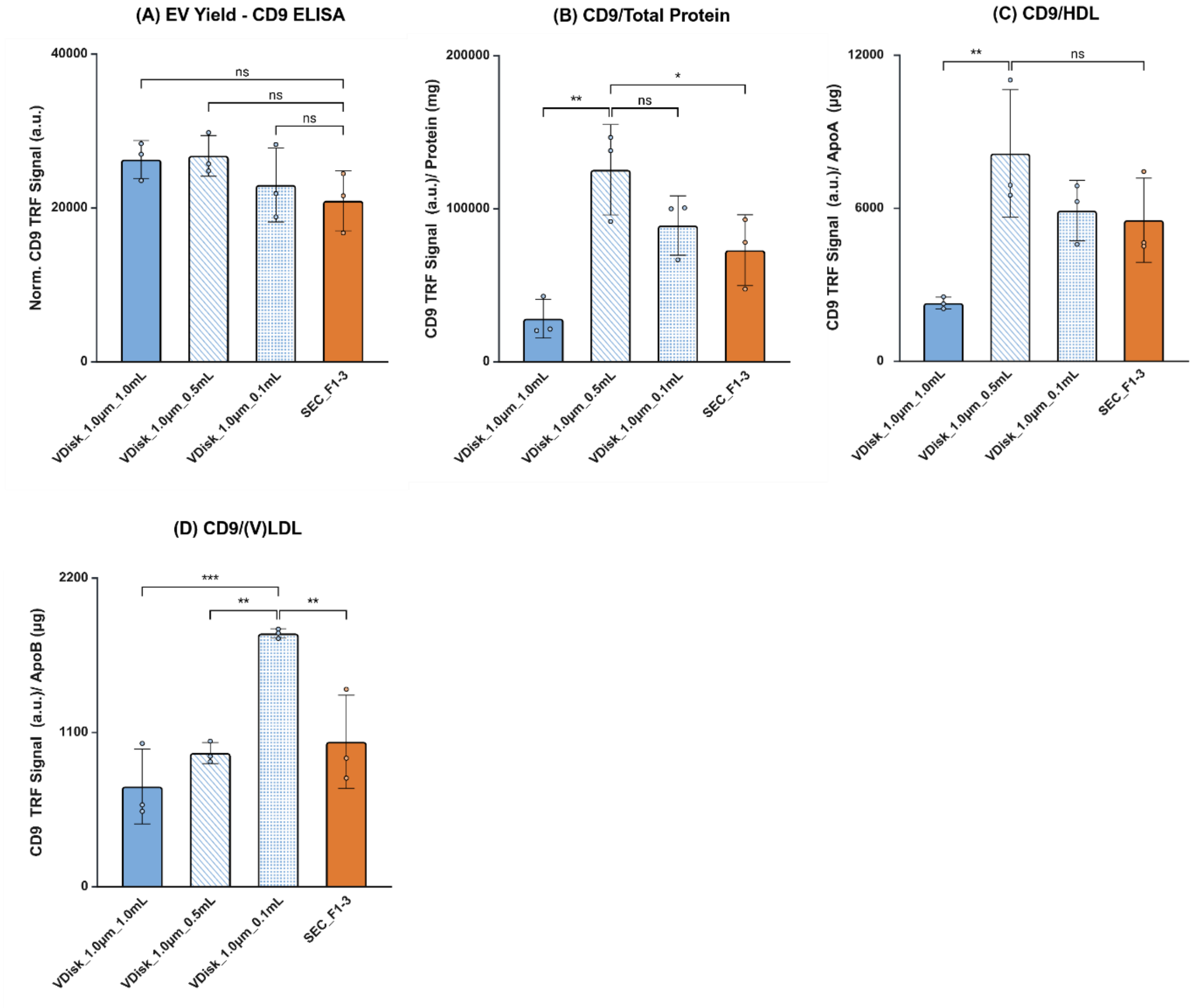


***Figure S6. VDisk_1.0µm performance at varying plasma volumes compared with SEC.*** *(A) EV yield (normalized CD9 TRF signal), (B) CD9 TRF signal per total protein, (C) CD9 TRF signal per ApoA (HDL marker), and (D) CD9 TRF signal per ApoB (V/LDL marker). Bars show means from three healthy donors (biological replicates); each point represents an individual donor. Error bars indicate standard deviations. Statistical analysis: one-way ANOVA with Tukey's multiple comparisons test (n = 3). Significance levels: ns = not significant, *p < 0.05, **p < 0.01, ***p < 0.001. Created with BioRender.com.*

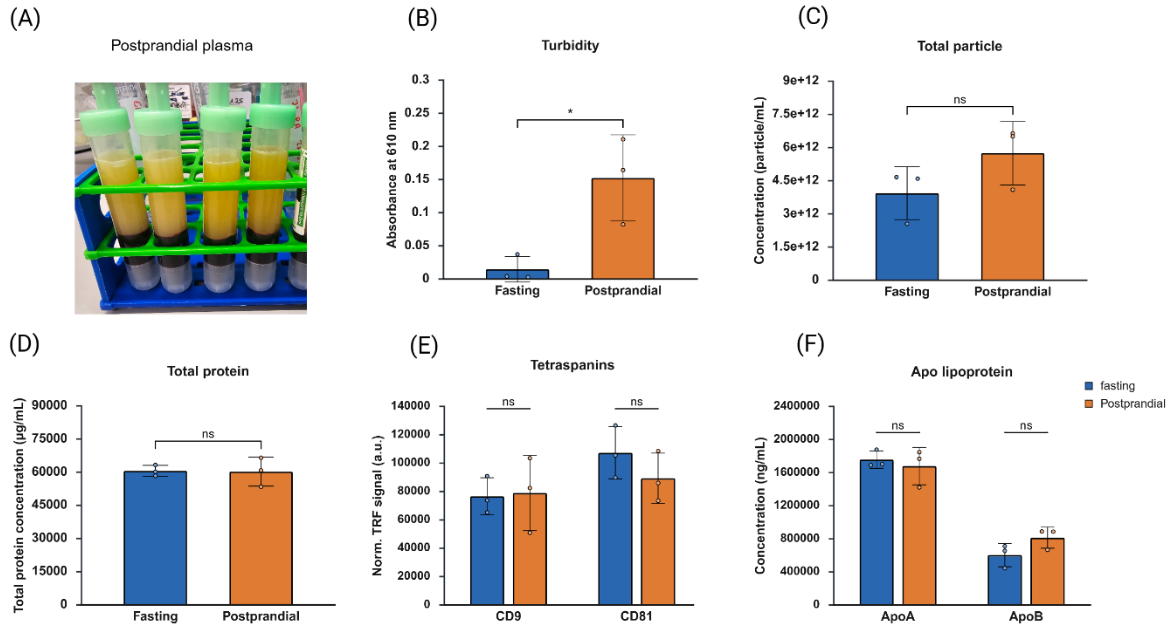


***Figure S7. Initial plasma matrix characterization****. (****A****) Appearance of postprandial plasma after the first centrifugation step. (****B****) Turbidity of 1:10 diluted plasma samples after plasma separation. (****C****) Total particle concentration measured by NTA. (****D****) Total protein content measured by the BCA assay. (****E****) Total CD9 and CD81 signal measured by ELISA. (****F****) Total ApoA concentration (HDL marker) measured by ELISA. In (****B–F****), each bar represents the mean of biological replicates (n = 3 healthy donors); each data point corresponds to the average of three technical replicates, and error bars indicate standard deviations. Statistical analysis: two-way ANOVA with Tukey's multiple comparisons test (n = 3). Significance levels: ns = not significant, *p < 0.05, **p < 0.01, ***p < 0.001. Pre-isolation measurements—particularly for NTA and ELISA—should be interpreted with caution, as high levels of contaminating proteins and lipoproteins in complex biological matrices may lead to apparent concentrations that do not accurately reflect true EV abundance. Created with BioRender.com.*

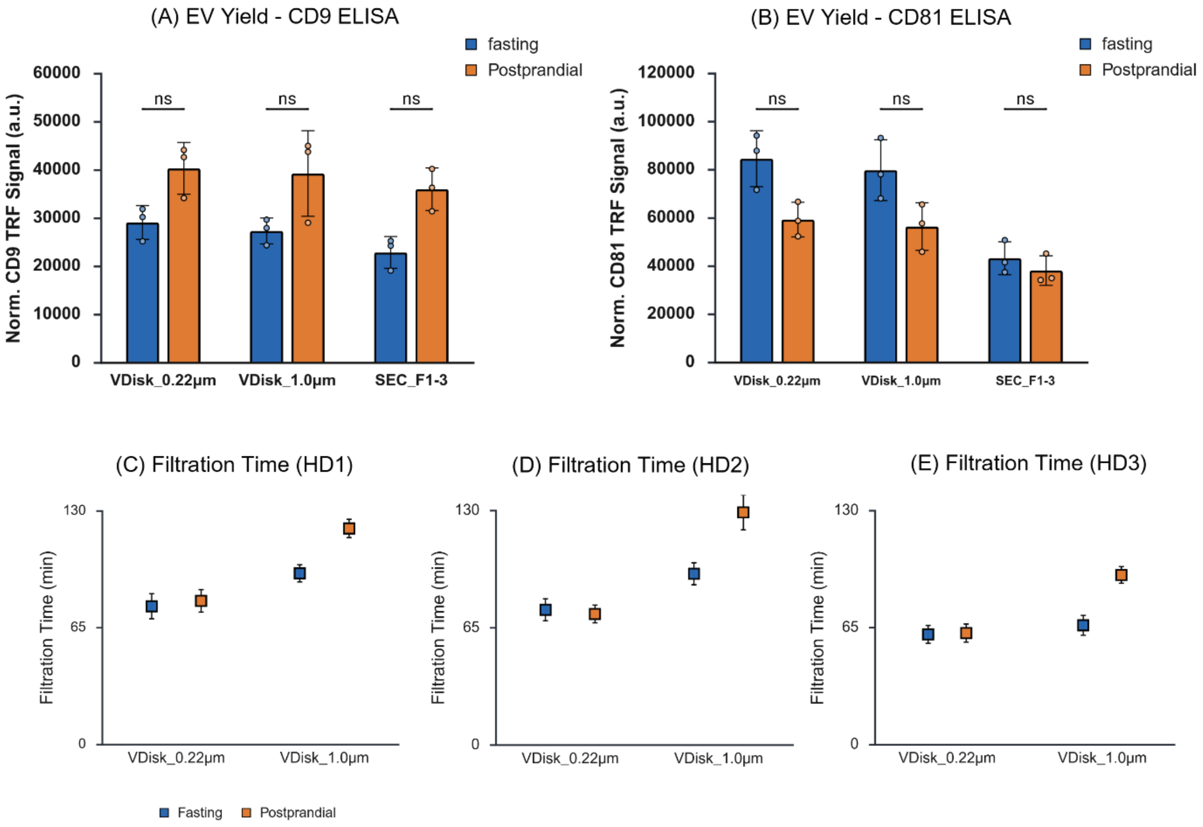


***Figure S8. EV yield and filtration times across different isolation methods.*** *(A–B) Normalized EV yield of VDisk_0.22µm, VDisk_1.0µm, and SEC_F1–3 based on CD9 and CD81 ELISAs in fasting and postprandial plasma. Each bar represents the mean of biological replicates (n = 3 healthy donors); each data point corresponds to the average of three technical replicates, and error bars indicate standard deviations. Statistical analysis: two-way ANOVA with Tukey's multiple comparisons test (n = 3). Significance levels: ns = not significant, *p < 0.05, **p < 0.01, ***p < 0.001. (C–E) Filtration times of VDisk_0.22µm and VDisk_1.0µm processing 1 mL of fasting or postprandial plasma. Created with BioRender.com.*

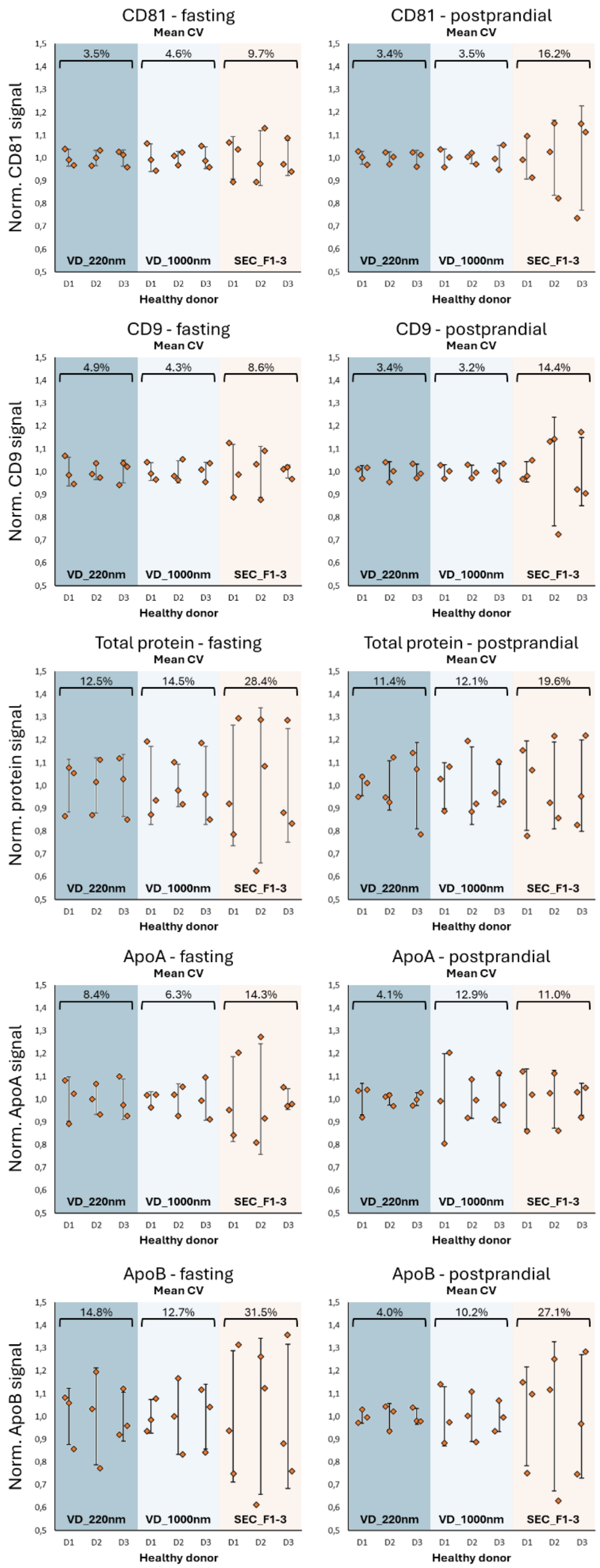


*Figure S9: Intra donor extraction variability. Normalized signals (CD81, CD9, total protein, ApoA, ApoB) of Disk and SEC extracted samples of three healthy donors (D1-D3). Each Sample was extracted in three replicates. Each dot represents either one disk or one SEC column. Intra-donor CVs were calculated and mean CVs for each extraction methods are shown as an indicator of the reproducibility.*

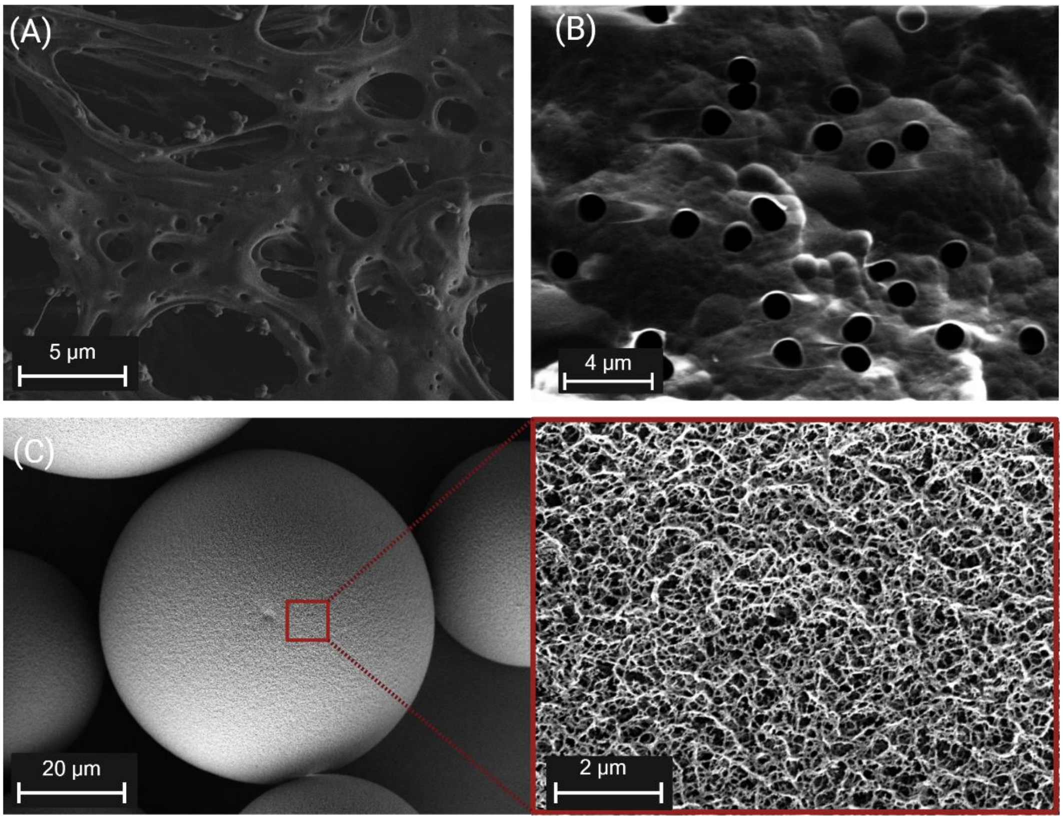


***Figure S10. Scanning electron microscopy (SEM) images.*** *(A) polyethersulfone (PES) membrane structure, (B) Polycarbonate track-etched (PCTE) membrane structure, and (C) surface morphology of MMC resin (CaptoCore 400 kDa).*

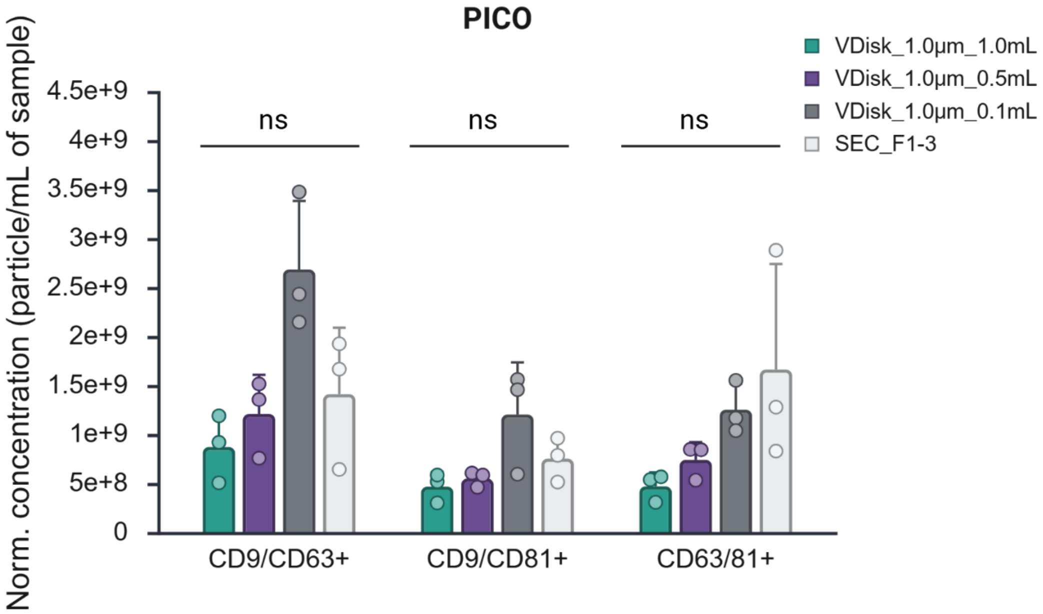


***Figure S11. PICO analysis of double-positive EVs.*** *Bars show means from three healthy donors (biological replicates); each point represents an individual donor. Error bars indicate standard deviations. Statistical analysis: two-way ANOVA with Tukey's multiple comparisons test (n = 3). Significance levels: ns = not significant, *p < 0.05, **p < 0.01, ***p < 0.001. Created with BioRender.com.*

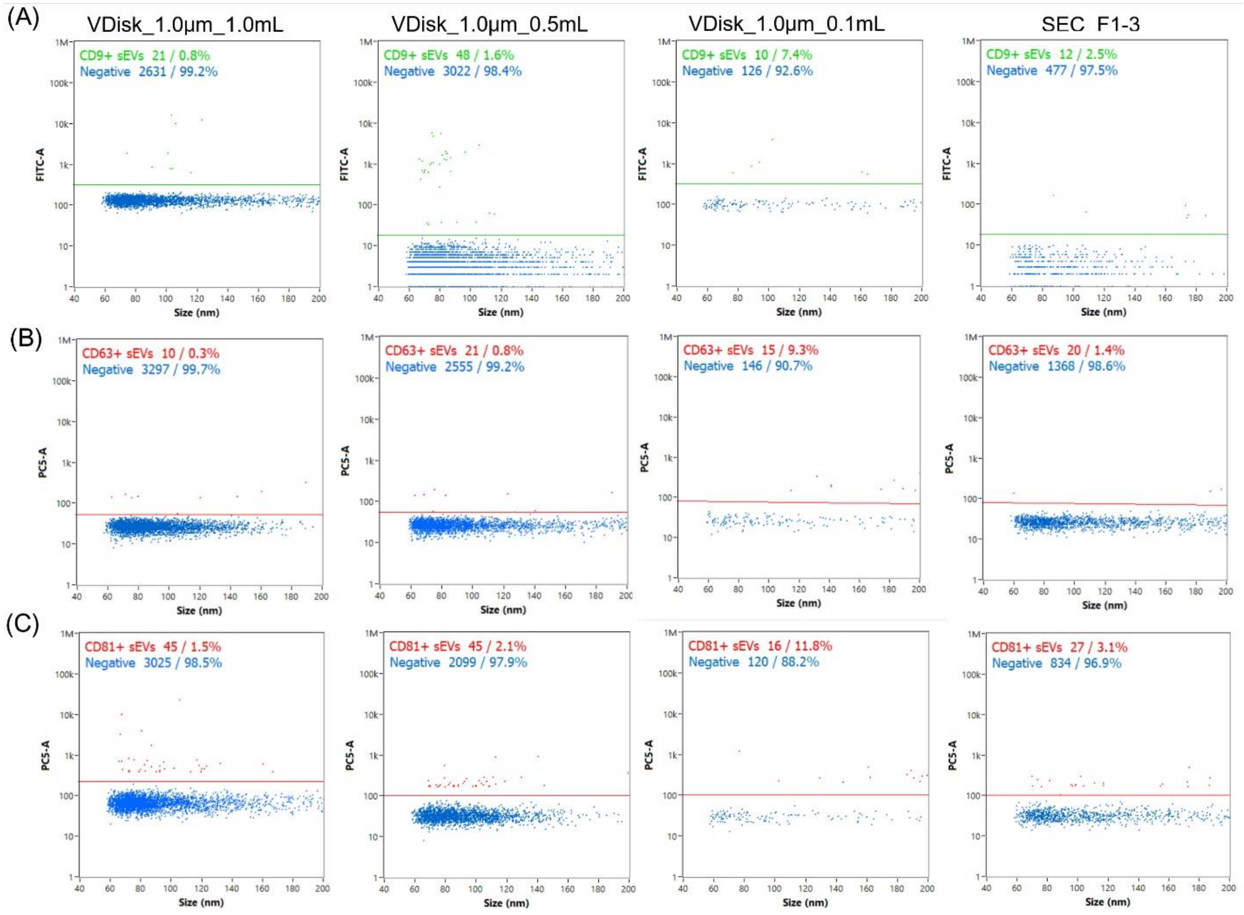


***Figure S12. NanoFCM analysis of EV surface marker expression across different isolation methods.*** *(**A**) $CD9^+$ and $CD63^+$ EVs, and (**B**) $CD81^+$ EVs measured in isolates obtained from VDisk_1.0μm using different plasma input volumes (1.0, 0.5, and 0.1 mL) and from SEC_F1–3. Shown is one representative replicate from three healthy donors (n = 3), incubated with antibodies against CD9, CD63, or CD81. Population 1 (P1; green/red) indicates marker-positive particles, whereas Population 2 (P2; blue) indicates marker-negative particles.*

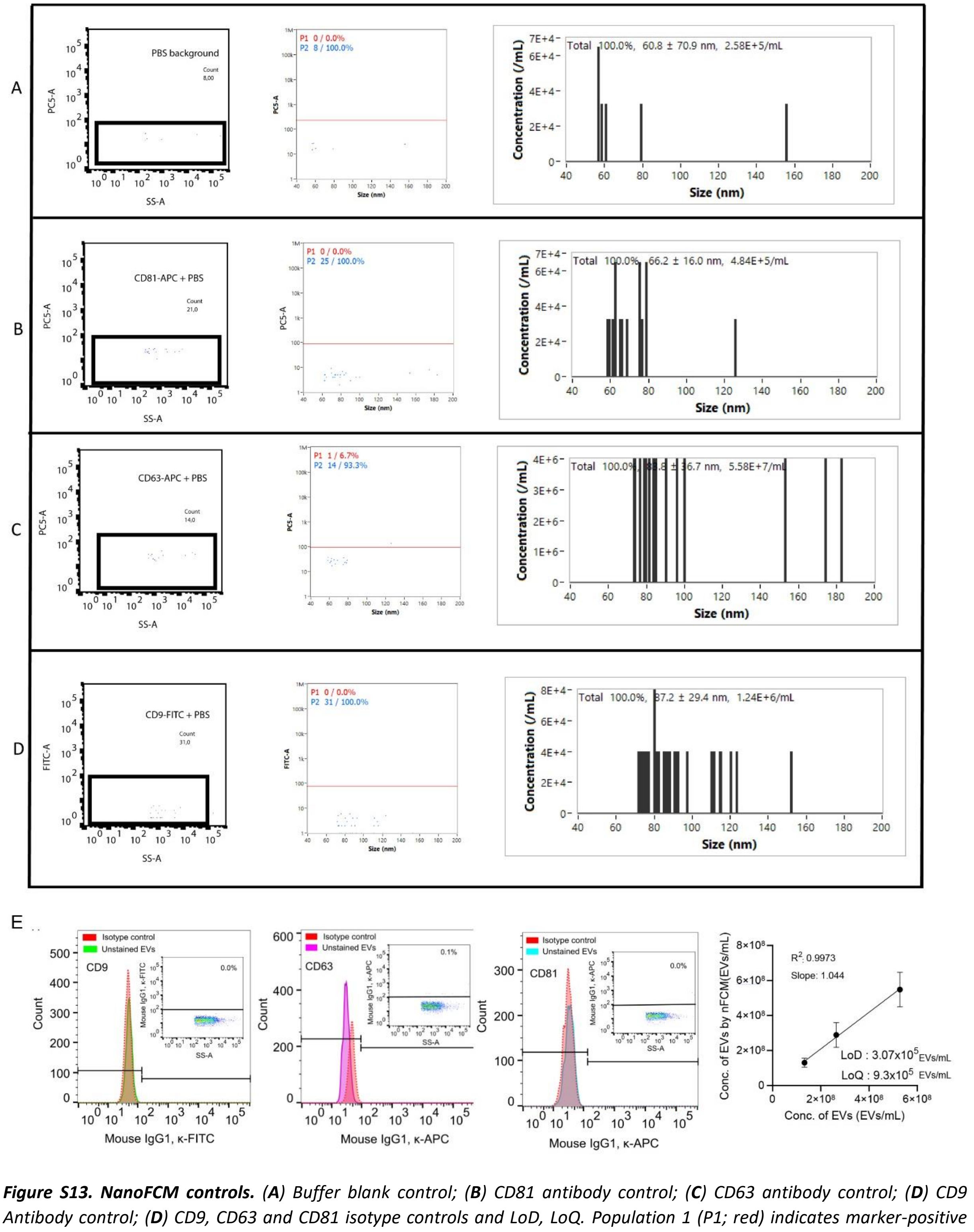

***Figure S13. NanoFCM controls.*** *(**A**) Buffer blank control; (**B**) CD81 antibody control; (**C**) CD63 antibody control; (**D**) CD9 Antibody control; (**D**) CD9, CD63 and CD81 isotype controls and LoD, LoQ. Population 1 (P1; red) indicates marker-positive particles, whereas Population 2 (P2; blue) indicates marker-negative particles.*

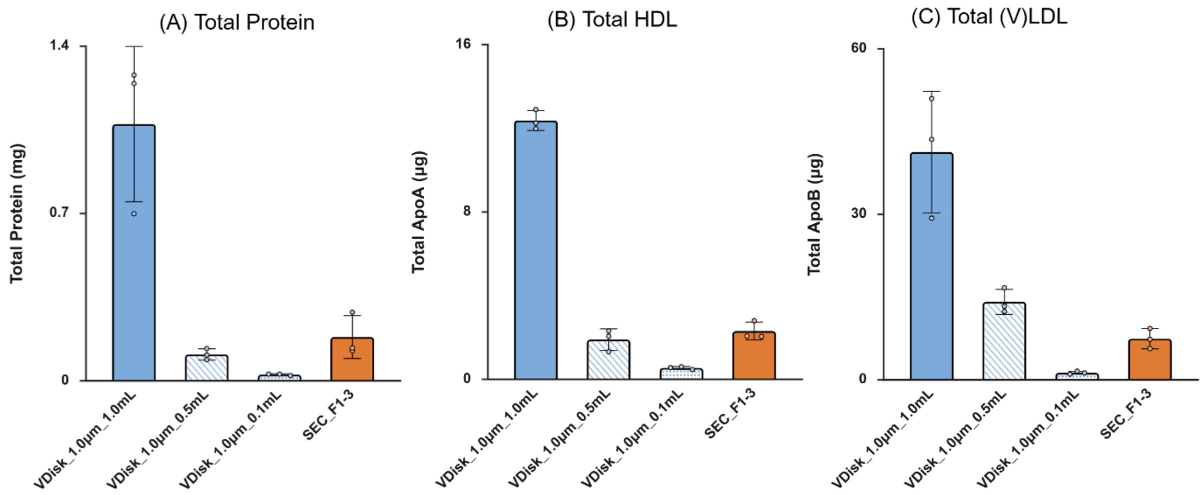


***Figure S14**. **Comparison of contaminant levels in EV isolates obtained from different VDisk configurations and SEC.** (A) Total protein, (B) total ApoA, and (C) total ApoB. Bars represent the mean values from biological replicates (n = 3 healthy donors), with individual data points displayed. Created with BioRender.com.*

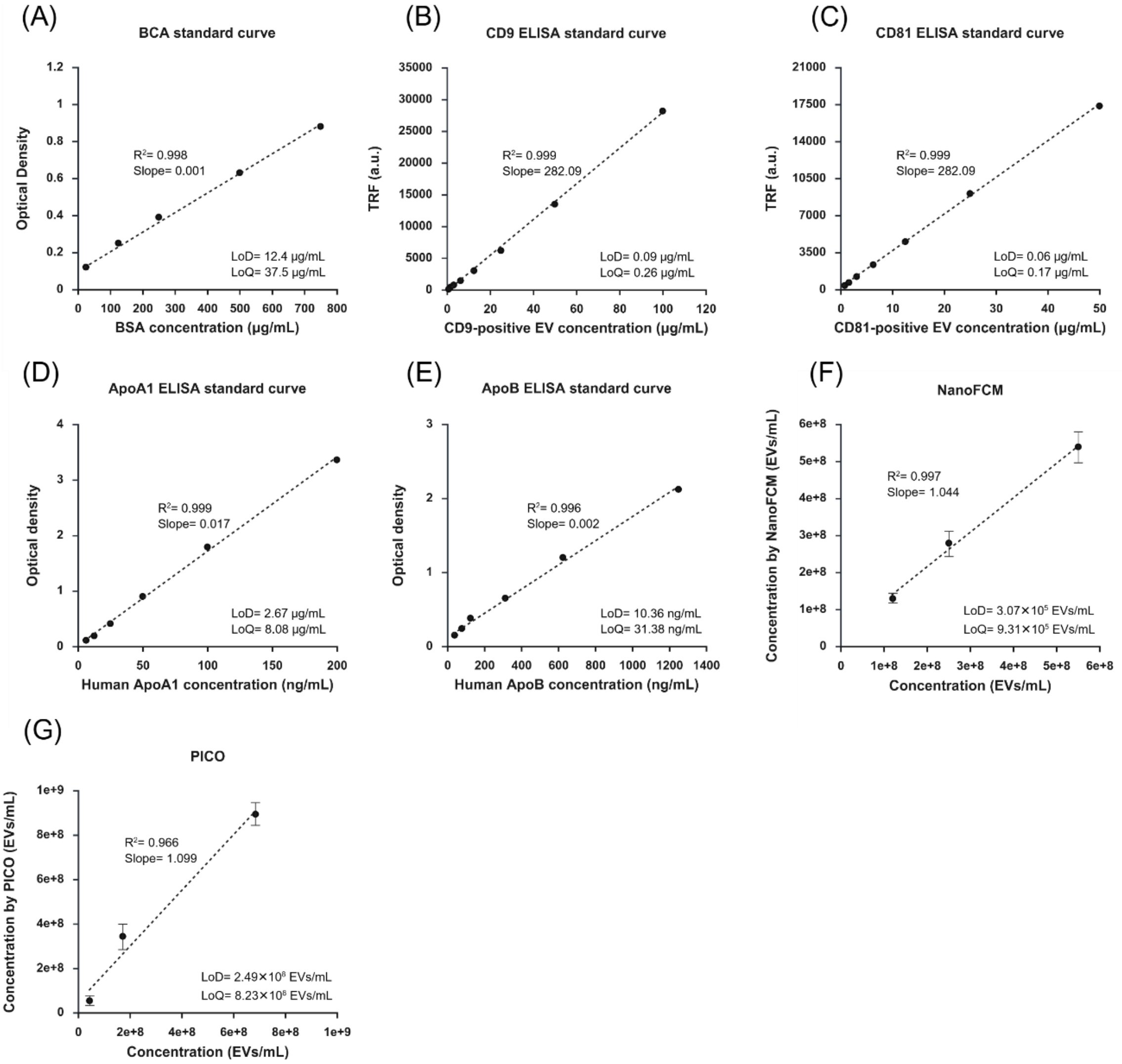


***Figure S15.*** *Standard curves for all analytical assays used in this study, including the limits of detection (LoD) and limits of quantification (LoQ) for each measurement. All sample concentrations were adjusted to fall within the lower limit of quantification of the respective assays. LoD and LoQ were calculated based on the calibration curve using the equations: LoD = 3.3×(σ/S) and LoQ = 10×(σ/S), where σ represents the standard deviation of the blank signal and S is the slope of the calibration curve.*
